# Benchmarking of shotgun sequencing depth highlights limitations of strain-level analysis and shallow metagenomics

**DOI:** 10.1101/2025.03.27.645659

**Authors:** Nicole S. Treichel, Charlie Pauvert, Joana Séneca, Petra Pjevac, David Berry, John Penders, Thomas C. A. Hitch, Thomas Clavel

## Abstract

Shallow metagenomics promises taxonomic and functional insights into samples at an affordable price. To determine the depth of sequencing required for specific analysis, benchmarking is required using defined microbial communities. We used complex mixtures of DNA from cultured gut bacteria and analysed taxonomic composition, strain-level resolution, and functional profiles at up to eleven sequencing depths (0.1-50.0 Gb). Reference-based analysis provided accurate taxonomic, and strain-level insights at 0.5-1.0 Gb. In contrast, *de-novo* metagenome-assembled genome (MAG) reconstruction required deep sequencing (>10 Gb), and even high-quality MAGs were chimeric, with 54.5 to 81.8 % accurately representing the original strains, depending on the bioinformatic approach used. However, the issue of chimeric MAGs can be reduced by using strain-aware assembly methods or long-read sequencing. Functionally, 2 Gb provided reliable insights at the pathway level, but sufficient proteome coverage was only achieved at or above 10 Gb. Library preparation and host DNA contamination were identified as confounders in shallow metagenomic analysis. This comprehensive analysis using complex mock communities provides guidance to an increasing community of scientists interested in using shallow metagenomics, and highlights the limitations of MAGs in accurately capturing strain-level diversity.

## Introduction

Next-generation sequencing is essential for microbiome research, enabling comprehensive analysis of complex microbial communities at an ever-increasing sample size. However, the widespread use of this technology contrasts with the low number of studies that have tested the robustness and accuracy of conclusions derived from widely used analysis workflows.

The most frequent used method for microbiota analysis is 16S rRNA gene amplicon sequencing due to its ease of implementation, both experimentally and bioinformatically, and low cost compared to metagenomics. However, it is limited to structural analysis of the microbiota (diversity and composition) and taxonomic resolution is generally limited to the genus level (strain analysis may be possible for certain taxonomic groups when using amplicon sequence variants). Experimentally it is also prone to amplification bias^1^. Whilst deep metagenomics offers a solution to these shortcomings, it is challenging to perform with low biomass samples or those with a high fraction of non-microbial DNA (e.g., human tissue samples) and is also more cost intensive.

Shallow metagenomics, commonly referred to as shotgun sequencing at a depth ≤1 Gb (ca. 3 million reads), has been proposed to overcome the limitations of both high-throughput 16S rRNA gene amplicon and deep metagenomic sequencing^2–4^. Few studies have attempted to benchmark shallow metagenomics and thereby define which information is accessible at what sequencing depth, and how reliably it can be obtained. A common approach used by most benchmarking studies is to compare sequencing techniques and depths using native samples, such as human stool^3–7^. The main disadvantage of this is that the exact composition of the input is unknown, preventing accurate benchmarking. Even fewer studies investigated the impact of sequencing depth using samples with defined microbial composition, hereon referred to as mock communities^2,8^. Cattonaro et al. used a commercially available mock community, containing 20 strains, of which the abundance of 5 was below their detection threshold^8^. Xu et al. used mock communities with two different distributions, including up to 69 human gut bacterial species^2^. However, their analysis relied on a limited set of sequencing depths and focused on comparing shallow metagenomics with 16S rRNA gene amplicon sequencing. Other common practices are to bioinformatically subsample deeply sequenced datasets to mimic different sequencing depths or to generate artificial mock communities in-silico^3,6,9^. However, these approaches do not include the impact of real sample processing and sequencing in the laboratory.

Given the need to better define the strengths and limitations of shallow metagenomics, we systematically evaluated the effects of shotgun sequencing depth on composition, strain-level diversity, and functional readouts (**Supplementary Fig. S1**). For this, we used 13 complex mixtures of bacterial DNA: an even distribution with 70 bacterial strains (Mock-even-70), a staggered distribution with 24 strains (Mock-stag-24), including 10 additional versions with different compositions (Mock-stag-24 v1-10), or a staggered distribution with 70 strains (Mock-stag-70) (**Supplementary Table S1**, **Supplementary Fig. S2**). As wet-lab factors can cofound results, DNA libraries were prepared in two laboratories and in the presence/absence of background DNA isolated from the gut of germfree mice.

## Methods

### DNA extraction and preparation of the Mock communities

For the creation of the mock communities (Mock-even-70, Mock-stag-24, Mock-stag-24 v1-10 for multi-coverage binning, and Mock-stag-70), DNA was obtained from isolates in our collections of mouse and human gut bacteria ^10,11^ (**Supplementary Fig. S2)**. DNA was extracted from freshly grown strains revived from frozen glycerol stocks based on the method of Godon et al. 1997 ^12^ modified as described in Afrizal et al ^10^. DNA concentration was measured using a Qubit fluorometer (Thermo Fisher Scientist). The DNA of all 70 isolates was pooled in either equimolar amounts (Mock-even-70), or in a staggered distribution (Mock-stag-70). For 24 selected isolates, the DNA was pooled in a staggered distribution (Mock-stag-24). For testing multi-coverage binning to generate MAGs, ten additional mock communities with 24 DNA extracts in 10 different distribution patterns were created (Mock-stag-24 v1-10). The distribution of DNA from the different isolates in each Mock community is provided in **Supplementary Table S1**.

Non-bacterial background DNA was isolated, as described above, using gut content collected from germfree mice (LAVE ethical approval 81-02.04.2023.A253) in accordance with the German Animal Protection Law (TierSchG). All mice were fed a standard chow *ad libitum* (γ-irradiated ssniff Spezialdiäten ref. V1124-927). The germfree status of the mice was confirmed by microscopic observation after Gram-staining and by cultivation on both anaerobic and aerobic agar plates. Mice were culled and caecal content was collected and stored immediately at -80°C. After DNA extraction, the background DNA was mixed 1:1 (v/v) with Mock-even-70 and Mock-stag-24.

### Library preparation and short-read sequencing

In sequencing facility 1 (UKA), DNA libraries of the mock communities were prepared using the NEBNext Ultra II FS DNA Library Prep Kit for Illumina (NEB, USA) according to the manufacturer’s instructions, using 100 ng (Mock-even-70 and Mock-stag-24), 200ng (Mock-stag-70) or 40 ng (Mock-stag-24 v1-10 for multi coverage binning) of input DNA and an automated platform (Beckman Coulter, USA). Enzymatic shearing to approximate 250 bp was performed for 30 minutes. Adaptor-ligated DNA was enriched using PCR (Mock-even-70, Mock-stag-24, Mock-stag-70: 5 cycles; Mock-stag-24- v1-10 multi-coverage binning: 7 cycles) and NEBNext Multiplex Oligos for Illumina (NEB, USA) for unique dual barcoding. AMPure beads (Beckman Coulter, USA) were used for size selection and clean-up of adaptor ligated DNA.

Sequencing facility 2 (UMC) used the Nextera XT DNA Library preparation kit (Illumina, USA) with Nextera XT indexes (Illumina, USA) and 1 ng template DNA according to manufacturer’s protocol. Twelve cycles of PCR were used for indexing. AMPure beads (Beckman Coulter, USA) were used for double-sided selection and clean-up of adaptor ligated DNA.

For cleaned DNA library from both facilities fragment size (∼320 bp library fragment size ≙ ∼200 bp genomic DNA insert size) was determined on an Agilent D1000 Tapestation (Bioanalyzer System, Agilent Technologies, USA) using High Sensitivity D1000 screentapes.

Quality check (Bioanalyzer System, Agilent Technologies, USA), DNA quantification (Quantus, Promega, USA), and sequencing of the resulting libraries were conducted at the IZKF Core Facility Genomics (UKA, RWTH Aachen University). The libraries for Mock-even-70, Mock-stag-70 and Mock-stag-24 were pooled to reflect the nine or eleven different sequencing depths targeted, and sequenced on a NovaSeq6000 (Illumina, USA) with NovaSeq 6000 Reagents v1.5 (2x150 cycles). Mock-stag-24 v1-10 for multi-coverage binning were sequenced at 10 Gb with the same chemistry at the NGS Competence Center Tübingen (NCCT).

### Long-read sequencing

The Mock-stag-70 DNA (200 fmol) was prepared for long read sequencing with the SQK-LSK114 kit (Oxford Nanopore Technologies), following the manufacturer’s protocol and the NEBNext Companion module v2 (New England Biomedicals). About 80 fmol of a >20Kb library was loaded on a R10.4.1 flowcell (FLO-PRO114, Oxford Nanopore Technologies) and sequenced for 72h on a Promethion P2 solo (Oxford Nanopore Technologies) using Minknow v. 25.05.12. The flowcell was washed using the EXP-WSH004 kit and leftover library was re-loaded and left to sequence until the flowcell end of life. Reads were base called using Dorado (v. 1.0.0) using super accuracy mode (model: r1041_e82_400bps_sup_v5.2.0). Long read sequencing was performed at the Joint Microbiome Facility of the Medical University of Vienna and the University of Vienna under project ID JMF-2507-23.

### Bioinformatic analysis

An overview of the bioinformatic workflow is provided in **Supplementary Fig. S3**. All steps are described in detail in the following sections.

### Reference genomes

The genomes of all strains used in the Mock communities are hereon referred to as “reference genomes”. They have been deposited in a public repository and published previously^10,11^. A phylogenetic tree was constructed from the genomes of the 70 isolates using PhyloPhlAn^13^ v3.0.67 (options: *--diversity medium –f supermatrix_aa.cfg*) (**Supplementary Fig. S2**). Genome characteristics, including size and GC content, were analysed using Biopython^14^ v.1.79 and bioawk^15^ v1.0 (**Supplementary Table S2**).

### Pre-processing of shotgun metagenomic data

Raw reads of samples with the different empiric sequencing depths (**Supplementary Table S3**) were further subsampled bioinformatically to the exact targeted sequencing depth (below). The raw FASTQ files were subsampled using seqtk^16^ v1.2 with default settings. The targeted number of paired-end reads (2 x 150 bp) per sequencing depth was as follows: 333,333 read pairs (0.10 Gb), 833,333 (0.25 Gb), 1,666,667 (0.5 Gb), 2,500,000 (0.75 Gb), 3,333,333 (1.0 Gb), 5,000,000 (1.5 Gb), 6,666,667 (2.0 Gb), 16,666,667 (5.0 Gb), 33,333,333 (10.0 Gb). Adapters were removed and subsampled raw reads were quality-filtered with Trimmomatic^17^ v.0.39 (options: *TRAILING:3 LEADING:3 SLIDINGWINDOW:5:20 MINLEN:50 ILLUMINACLIP:{adapters.fa}:2:30:10*). The bbduk command (options: *hdist=1 k=31*) in BBMap^18^ v.38.84 was used for removal of phiX sequences. Quality-filtered reads of all Mock communities were assembled into contigs using MEGAHIT^19^ v1.2.9. Additionally, the quality-filtered reads of Mock-stag-70 were assembled into contigs with metaSPAdes v.4.2.0^20^. Contigs of metagenomes sequenced at 10 Gb, 20 Gb and 50 Gb assembled with either of the two assemblers were aligned to the reference genomes with blastn (perc_identity, 97%; evalue, 1e-10; alignment length, >150 bp). In addition to the 3 mock communities sequenced separately at all sequencing depths, and to the 10 Mock-stag-24 communities with different compositions (v1-10) sequenced at 10 GB, a further validation step included in-silico subsampling of the 50 Gb Mock-stag-70 to 10 sequencing depth (0.1 Gb – 20 Gb), each ten times, using seqtk^16^ v1.2 (default settings with 10 different seeds).

### Taxonomic coverage

Coverage of the reference genomes by the quality-filtered metagenomic reads was determined using coverM^21^ v0.6.1 (options: *coverm genome --mapper bwa-mem --methods covered_fraction --min-covered-fraction 0 --coupled*). The relative abundance profiles were calculated using the read count option of coverM (options: *coverm genome --methods count --min-covered-fraction 0 --coupled*), which were converted to relative abundance for each sample.

Non-supervised taxonomic profiles were generated using MetaPhlAn v4.0.1^22^ with the mpa_vJun23_CHOCOPhlAnSGB_202403 database.

Average nucleotide identity (ANI) values between the reference genomes of four *Escherichia coli* strains and four *Phocaeicola vulgatus* strains were calculated using FastANI^23^ v1.34.

### Functional analysis

Protein-coding genes were predicted in the pooled reference genomes and the assembled contigs of the Mock samples (individually) using prodigal ^24^ v2.6.3.

Gene functions were annotated using KEGG Orthology and KofamScan^25^ v1.3.0. Pathway completeness was assessed by KEGG-Decoder^26^ v1.3, as percentage of KOs covered, which are included in the manually curated canonical pathways used by KEGG-Decoder.

To determine the ability of each sequencing depth to capture the protein-encoding potential in the Mock communities, the genes predicted in the reference genomes were used to create a Diamond^27^ (v2.0.15) protein sequence database, with which the protein sequences predicted in the assemblies (one assembly for each of the sequencing depths) were compared (options: *diamond blastp –sensitive –query-cover 80 –id 90*).

### Metagenome-assembled genomes (MAGs)

Contigs <1,000 bp were removed prior to reconstructing MAGs. An index table was built from the size-filtered contigs using bowtie2^28^ v2.5.1 (*bowtie2-build*) with default options. The decontaminated paired-end reads were aligned to the bowtie index of size-filtered contigs (*bowtie2 –S - --very-sensitive-local –no-unal –p 30*). Bam files were sorted with samtools^29^ v1.17 (*samtools view –bS).* Contigs were binned using Metabat2^30^ v2.12.1 and its algorithm for calculating coverage of each sequence in the assembly (*jgi_summarize_bam_contig_depths*) before creating the bins (*metabat2 -m 1500 --maxP 95 --minS 60 --maxEdges 200 --unbinned --seed 0*).

The quality of the resulting bins was evaluated with checkM^31^ v1.1.3 using the lineage workflow. Accordingly, the MAGs were categorised as being of high quality (>90% completeness, <5% contamination), medium quality (>70% completeness, <10% contamination), or low quality (all that did not fulfil the previous criteria). They were then taxonomically classified using GTDB-Tk^32^ v2.3.2 (*gtdbtk classify_wf*) with the Genome Taxonomy Database r207. The coverage of the reference genomes by the different MAGs was determined using coverM^21^ v0.6.1 (*coverm genome --mapper bwa-mem --methods covered_fraction --min-covered-fraction 0 –single; no multiple read mapping*). Reference genomes were aligned to MAGs by blastn v2.13.0 *(-evalue 1e-10 -perc_identity 90.0*) and circular alignments plotted using shinyCircos^33^.

For comparing sample-specific (approach above) to multi-coverage binning for the ability to reconstruct high-quality, non-chimeric MAGs (i.e. one genome match only), the workflow of Mattock and Watson^34^ was used on Mock-stag-24 v1-10, sequenced at a depth of 10 Gb each.

### Long-read sequence analysis

The raw reads were filtered to a minimum average read quality of Q20 and a minimum length of 1,000 bp using chopper (v0.10.0)^35^. The filtered reads were used to randomly subset 11 additional datasets corresponding 0.1, 0.25, 0.5, 0.75, 1, 1.5,2, 5, 10, 20, and 50 Gb using rasusa (v2.1.1)^36^. Each subset was subsequently assembled using flye (v2.9.5)^37^ with “–nano-hq” and polished once with medaka (v2.1.0, github.com/nanoporetech/medaka) using the –bacteria flag. Contigs <1,000bp were removed. Assemblies were quality checked using QUAST^38^ v5.3.0 and metagenomic binning was performed using metabat2^30^. Bin coverage was estimated using the *jgi_summarize_bam_contig_depths* function with a minimum percent identity of 90. Quality and chimeric nature of MAGs was assessed as described above for the short-read data.

### Statistics and Plotting

All statistical tests were performed in R^39^ v4.4.1. using RStudio v2023.03.0+386 and the packages vegan^40^, reshape^41^, tidyverse^42^ and dplyr^43^. For creation of the graphs, the R package ggplot2^44^ v3.5.1, ggpubr^45^ v0.6.0, ggbreak^46^ and ggfortify^47^ v0.4.17 were used. The specific statistical tests used in each analysis are stated in the results.

## Results

### Shallow metagenomics enables reliable reference-based taxonomic composition

We first investigated the impact of sequencing depth on taxonomic composition. For the three main Mock communities (Mock-even-70, Mock-stag-24, Mock-stag-70), reads were detected for all reference genomes already at 0.1 Gb. To obtain an overview of genome coverage, completeness categories were compared between the sequencing depths. At 0.1 Gb, most of the genomes (63 to 91 %) had a low coverage (0-25 %) (**Fig. 1A**). Genome coverage increased substantially until 5 Gb of sequencing, with a clear effect of mock community complexity. Most genomes reached >90 % coverage at 5 Gb in Mock-even-70 and Mock-stag-24. However, in Mock-stag-70, coverage continued to increase gradually from 5 Gb (36 % genomes at >90 % coverage) to 50 Gb (64 %), but 11 genomes (16 %) added with 0.001 ng (0.00046‰) to 0.1 ng (0.0046 %) had still low coverage (0-25 %) at the highest sequencing depth (50 Gb).

**Fig. 1:**
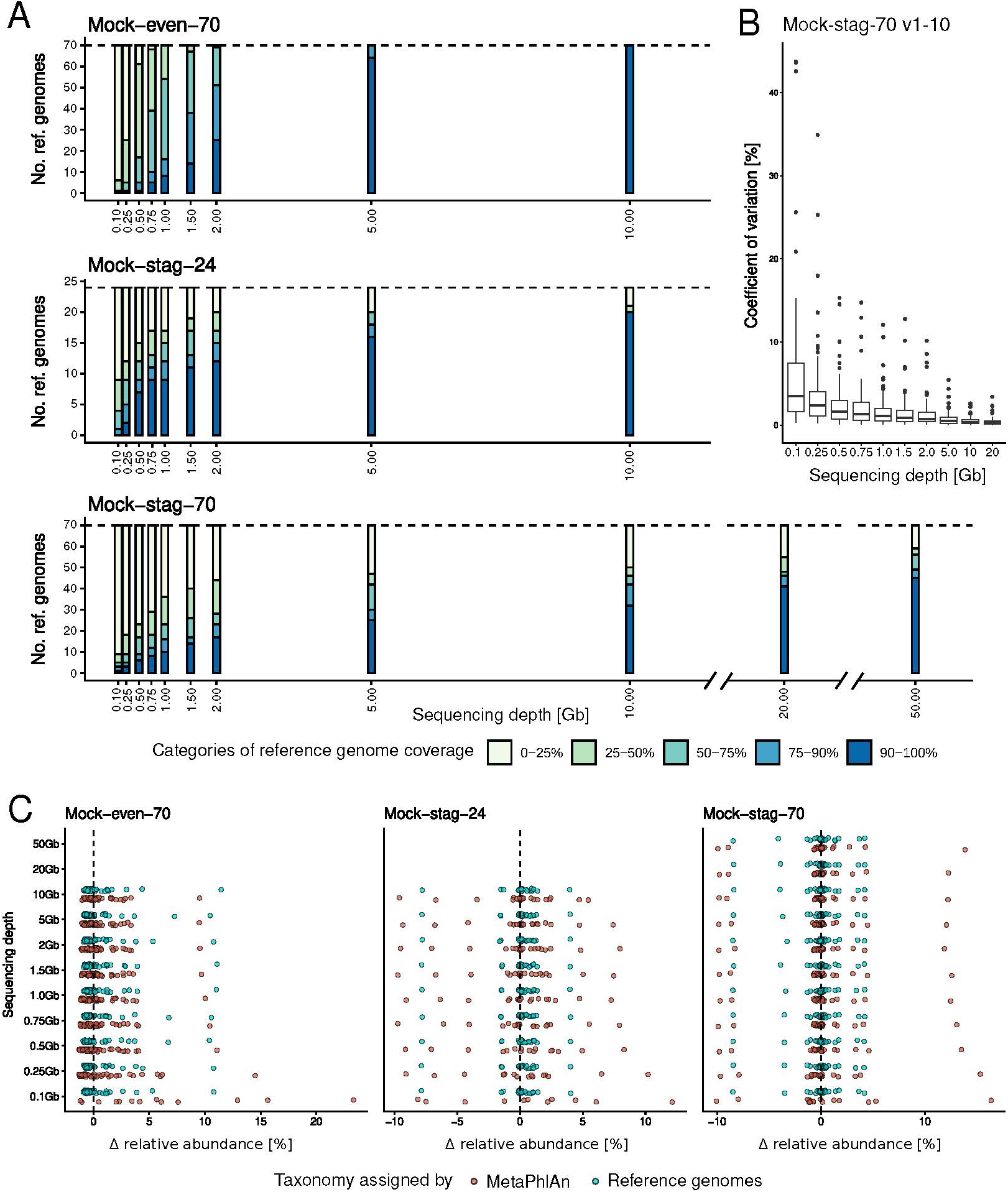
Reference-based taxonomic profiles. **A** Number of reference genomes per category of coverage by metagenomic reads (colour gradients) for Mock-even-70, Mock-stag-24, and Mock-stag-70 (top to bottom) at up to eleven sequencing depths (x-axis). **B** Coefficients of variation of relative abundances of the 70 strains in ten *in-silico* datasets for each sequencing depth, subsampled from Mock-stag-70 sequenced at 50Gb. The coefficients of variation between the sequencing depths were tested statistically using a Kruskal-Wallis rank-sum test with Benjamini-Hochberg correction (p-values in **Supplementary Table S5**). **C** Difference (delta-values) between measured relative abundances and theoretical values (x-axis) for the three mock communities after taxonomic assignment using either the reference genomes (blue) or MetaPhlAn4 (red). In the MetaPhlAn analysis, only the taxa that matched reference genomes at the species level were considered.

We then looked at the individual strains across the sequencing depths (**Supplementary Fig. S4**). In Mock-even-70, the lowest coverage at 0.1 Gb was 3.34 % for *Phocaeicola sartorii* CLA-AV-12 and the highest 58.7 % for *Bacteroides uniformis* CLA-AV-11, which reached 99 % coverage already at 0.75 Gb. Full genome coverage (100 %) was only achieved at 10 Gb for V*eillonella intestinalis* CLA-AV-13. In Mock-stag-24 and Mock-stag-70, higher coverage was associated with increasing DNA concentration. The reference genomes with lowest concentration in Mock-stag-24 (0.01 %, 0.04 ng) had a minimum coverage of 0.53 % at 0.1 Gb and a maximum coverage of 34.21 % at 10 Gb. In Mock-stag-70, the lowly abundant genomes - 0.1ng (0.0046%) to 0.001 ng (0.00046‰) - reached a coverage of 0.8 % to 21.8 % at highest sequencing depth (50 Gb), whilst the genomes of the five strains with highest DNA amounts (>4.6 %) reached between 40.50 % and 98.25 % coverage already at 0.1 Gb.

Next, we assessed the relative abundance of strains and compared them to their theoretical values. For Mock-even-70, relative abundances oscillated around the expected value of 1.43 % for most species, without any significant effects of the sequencing depth on the overall relative abundance profile (p-value = 0.78; Kruskall-Wallis test) (**Supplementary Fig. S5A**). Six species showed values >2.5 % for several sequencing depths, with highest values of 10-15 % for *B. uniformis* CLA-AV-11. In addition, marked differences due to sequencing depth were detected in the case of *Collinsella* sp. CLA-ER-H5. Sequencing depth had also no influence on the overall relative abundance profile in the case of Mock-stag-24 and Mock-stag-70 (p-value = 1 in both cases; Kruskall-Wallis) (**Supplementary Fig. S5B**). In addition to sequencing the mock communities separately, Mock-stag-70 sequenced at 50 Gb was subsampled multiple times (10 replicates) bioinformatically for each sequencing depth ≤20 Gb to test variability. Variations in the relative abundance of individual species was low in general (average coefficients of variation <5 %), with increasing variability at lower sequencing depths (**Fig. 1B**; statistics in **Supplementary Table S5**).

As reference genomes are not available for metagenomic analysis in most studies, the mock communities were also taxonomically analysed using the commonly used profiler MetaPhlAn^22^, hereon referred to as non-supervised approach. Overall, the taxonomic assignment in this approach was less sensitive, *i.e.*, fewer species were detected. In Mock-stag-24, one strain with low concentration (0.1 %, 0.4 ng, *Hominilimicola fabiformis* CLA-AA-H232) was not detected (**Supplementary Fig. S6**). In Mock-even-70, 59 species could be assigned with MetaPhlAn4, including 47 matching a reference genome at species level. In Mock-stag-70, 53 taxa were detected, 44 with a species-level match (**Supplementary Fig. S6**). The non-supervised taxonomic assignment showed also higher variance from the targeted relative abundances, especially at lower sequencing depths and with increased complexity of the mock communities (**Fig. 1 C**; statistics in **Supplementary Table S5**).

In summary, reference-based detection of strains is possible with sequencing as little as 0.5 Gb, but high coverage of reference genomes needs more data (>5 Gb) under the conditions tested. Relative abundance profiles were not substantially affected by sequencing depths for most of the strains. In contrast, non-supervised taxonomic assignment was less sensitive, with fewer species detected.

### Strain-level analysis: *De-novo* MAG reconstruction requires deep sequencing and generates chimeras

The ability of shallow metagenomics to resolve strain-level variation was investigated using four strains of *E. coli* and *P. vulgatus* within the Mock-even-70 dataset (Supplementary **Fig. S2**). Coverage of their reference genomes per sequencing depth, as determined during assignment of all reference genomes, revealed that all strains were detected already at 0.1 Gb (Supplementary **Fig. S7A**). All four reference genomes of both species reached >75 % coverage at 5 Gb and >98 % at 10 Gb. All *P. vulgatus* strains showed a similar increase in coverage with increasing sequencing depth, mirroring ANI values between the strains that were all close to one another (Supplementary **Fig. S7B**). In contrast, for the *E. coli* strains, the reference genomes sharing the highest ANI value (*E. coli* Ec8850 and CR B11; ANI = 99.97 %) showed a congruent coverage increase (Supplementary **Fig. S7B**; their coverage increase curves overlaid), whilst *E. coli* CLA-AD-1, characterized by an ANI value more distant to the other strains (< 97 %), reached >99 % reference genome coverage already at the 2 Gb sequencing depth (Supplementary **Fig. S7B**). Only one strain of *E. coli* and *P. vulgatus* was detected in Mock-even-70 and Mock-stag-70 using MetaPhlAn4 at any of the sequencing depths (Supplementary **Figure S6**).

Reference datasets are rarely available during metagenomic analysis, hence *de-novo* reconstruction of genomes is widely used to facilitate strain-level analysis. MAGs were thus assembled separately for each metagenomic sample. Overall, the number of MAGs recovered increased with higher sequencing depth for all mock communities (**Fig. 2A**, Supplementary **Fig. S8, left panels**). More MAGs than the actual number of reference genomes (dashed line) were formed at the highest sequencing depths, whilst not all reference genomes were represented by MAGs (**Fig. 2A**, Supplementary **Fig. S8, right panels)**. The number of MAGs formed per reference genome increased with the amount of data sequenced, with ≥2 MAGs observed for 22, 12 and 29 of the reference genomes across all sequencing depths in Mock-even-70, Mock-stag-24, and Mock-stag-70, respectively (**Supplementary Fig. S9**). This suggests that multiple MAGs are created per reference genome with greater sequencing depth, rather than coalescing into a single high-quality MAG.

**Fig. 2:**
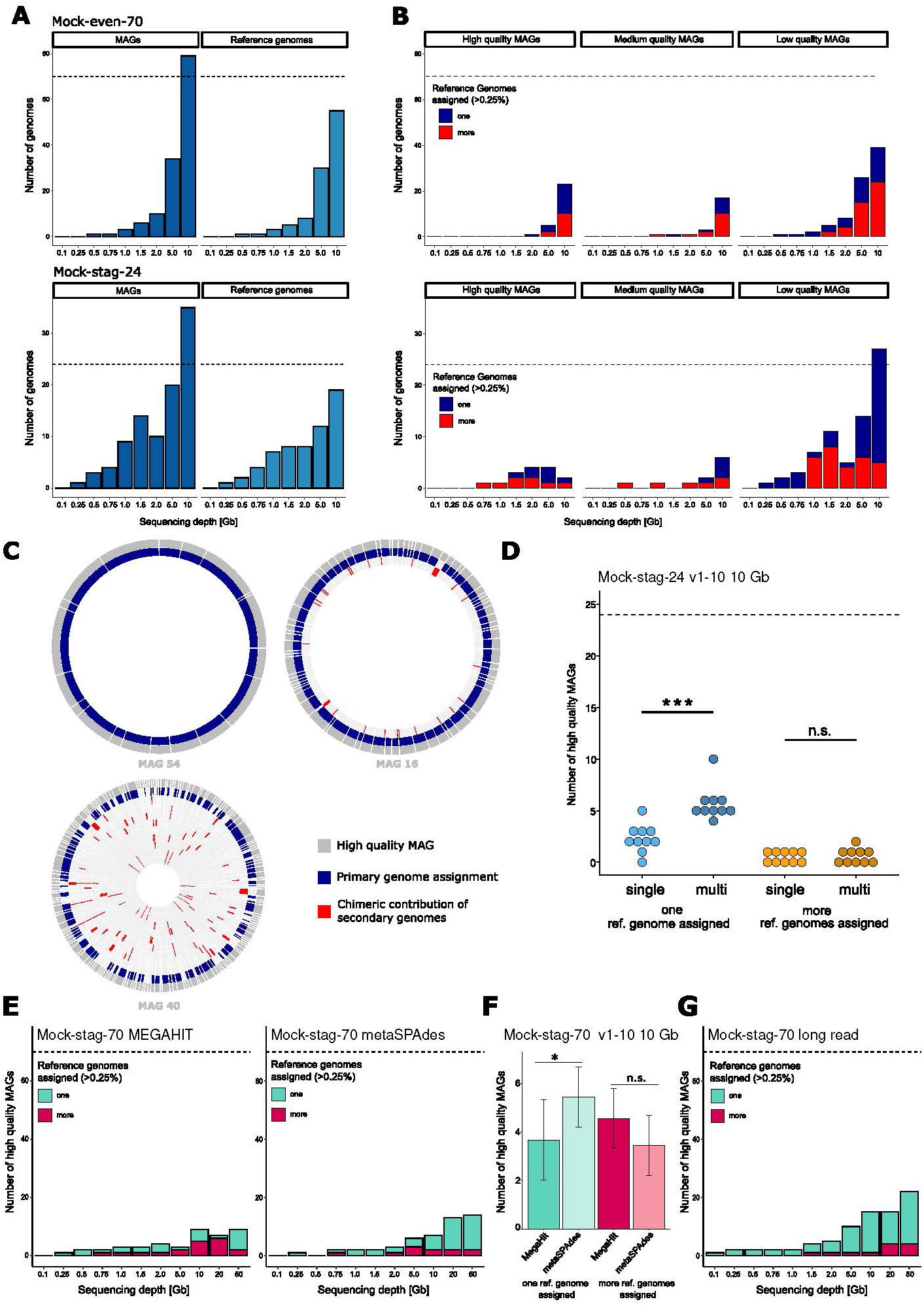
Strain analysis. **A** Strain analysis based on the MAGs assembled in Mock-even-70 (top) and Mock-stag-24 (bottom) at each sequencing depth individually. The left panels (dark blue) depict the number of bins assembled from the shotgun data; the right panels (lighter blue) show the number of reference genomes matching MAGs (coverM), indicating that multiple MAGs were reconstructed for some Mock species. The reference genome with the highest coverage by MAG reads was chosen as representative of that MAG. **B** Number of high-, medium- and low-quality MAGs assigned to either one (blue) or several (red) reference genomes (with coverM using a cut-off of > 0.25% coverage of reference genomes). **C** Three exemplary high-quality (> 90 % completeness, < 5% contamination) (hq) MAGs (dark grey, outer circle) reconstructed from Mock-even-70 at 10 Gb. They were aligned to the reference genomes they covered by more than 0.25% to illustrate different categories of chimerism. The predominantly covered reference genome is depicted in blue, while chimeric sequences of further reference genomes (inner circles) are shown in red. **D** Number of hqMAGs assigned to one reference genome (blue) or more reference genomes (orange) for the ten different Mock-stag-24 (v1-10) with varying reference genome abundance distribution. MAGs were constructed with either a single- (bright colours) or a multi-coverage (darker colours) binning approach. Statistics: Kruskal-Wallis rank-sum test; *** p-value = 0.0003871. **E** Number of hqMAGs constructed from contigs assembled with either MEGAHIT or metaSPAdes for Mock-stag-70. The MAGs were then assigned to one (green) or more (red) reference genomes as above. **F** High-quality MAGs binned from assemblies generated with either MEGAHIT or metaSPAdes using 10 datasets at 10 Gb subsampled *in-silico* from 50 Gb. Statistics: Wilcoxon rank-sum test; * p-value = 0,031. **G** Number of hqMAGs binned from assemblies acquired by long-read sequencing of Mock-stag-70 and categorised as in E.

Given the unexpected number of MAGs and their multiplicity per genome, we next determined their origin. MAGs were grouped by quality and their contigs were assigned to the reference genomes to determine which isolates they were reconstructed from, with a threshold of >0.25 % coverage of a reference genome. In Mock-even-70 at the highest sequencing depth (10 Gb), 10 of the 23 high-quality MAGs (hqMAGs; completeness >90 %, contamination <5 %) were assigned to multiple genomes (red bars) (**Fig. 2B, top**). The proportion of MAGs assigned to only one (blue) vs. multiple (red) genomes decreased further with decreasing quality of the MAGs. Taxonomic assignment of hqMAGs to GTDB showed a result congruent with the assignment to our own reference genomes (**Supplementary Table S4**). Although four *E. coli* strains were included in Mock-even-70, only one MAG assigned to *E. coli* CLA-AD-1 was reconstructed from each sequencing dataset > 1Gb, except 2 Gb (a second *E. coli* MAG of low completeness and quality was obtained). In the case of *P. vulgatus* (also represented by four strains), one MAG was found at 5 Gb and one at 10 Gb, both primarily assigned to *P. vulgatus* HDF, indicating strain delineation was not possible with the method used (**Supplementary Table S4**).

In Mock-stag-24, a single hqMAG was assembled at 0.75 and 1 Gb (**Fig. 2B, bottom**). Most sequences of this MAG were assigned to a genome with highest DNA concentration (40 ng, *T. ramosa* CLA-JM-H52, ≥95 % genome coverage). With increasing sequencing depth, some hqMAGs assigned to genomes with lower concentrations started to be reconstructed (**Supplementary Table S4**). Surprisingly, a higher proportion of single-origin MAGs occurred with decreasing quality, while at 10 Gb half of the hqMAGs in Mock-stag-24 were chimeric.

To illustrate the chimeric nature of MAGs, three exemplary hqMAGs (Mock-even-70, 10 Gb) were aligned to the reference genomes they covered >0.25 % using blastn (**Fig 2C**). MAG 54 was a best-case scenario with only one reference genome (*V. intestinalis* CLA−AV−13) matched with >98 % coverage. MAG 16 showed fragments of a second reference genome. MAG 40 represented a highly chimeric hqMAG, including 12 reference genomes (of which 10 are shown, ranked by decreasing coverage from outer to inner circle). The mean/maximum number of reference genomes per hqMAG at 10 Gb was 2.3/12, 1.5/2 and 2.1/5 for Mock-even-70, Mock-stag-24 and Mock-stag-70, respectively (**Supplementary Table S3**).

As multi-coverage binning was shown to increase the number and quality of MAGs^34^, we sequenced (10 Gb) ten additional Mock-stag-24 communities (v1-10) with varying reference genome distribution (Supplementary **Table S1**). In comparison to single-coverage binning, significantly more hqMAGs per mock community were assigned to only one reference genome using multi-coverage binning (**Fig. 2D**, blue dots). Despite this improvement, chimeric MAGs were still detected in half of the communities (**Fig. 2D**, orange dots).

To further assess ways to enhance MAG reconstruction, the most complex community (Mock-stag-70) was used to test the effects of assemblers (MEGAHIT – as in all analyses above – Vs. metaSPAdes) and the sequencing technology (short-read by Illumina- Vs. long-read by Nanopore-sequencing). MetaSPAdes assemblies generated more hqMAGs assigned to only on reference genome, especially ≥10 Gb (+1 at 10 Gb, +10 at 20 Gb, +5 at 50 Gb) (**Fig. 2E**). This was confirmed statistically by binning MAGs from assemblies generated with either of the two methods (MEGAHIT Vs. MetaSPAdes) using 10 datasets at 10 Gb subsampled in silico from 50 Gb (**Fig. 2F**). Long-read sequencing provided even more hqMAGs with a single reference match (14 of 15 at 10 Gb; 18 of 22 at 50 Gb) (**Fig. 2G**).

To know whether chimeric sequences are generated already during the assembly process, contigs were assembled from the 10 Gb, 20 Gb, and 50 Gb data using either MEGAHIT or metaSPAdes and were aligned to the reference genomes using blastn (Supplementary **Fig. S10**). With both assemblers, the majority of the contigs were assigned to only one reference species (average of three sequencing depths: MEGAHIT, 91.1 ± 1.6 %; metaSPAdes, 93.5 ± 1.0 %), suggesting that 6.5 to 8.9 % misassembled contigs contributed to the reconstruction of chimeric MAGs.

In summary, when high-quality reference datasets are available, read mapping facilitates strain-level analysis of shallow metagenomic data. *De-novo* analysis based on MAG reconstruction leads to contaminated genomes (chimeras), even at supposed high-quality. Chimerism stems from both the assembly and binning process, and was partially improved by multi-coverage binning for short reads, strain-aware assembly, or long-read sequencing. These findings suggest that strain-level diversity within MAG catalogues is artificially inflated due to chimerism. However, this issue can be reduced by use of strain-aware assembly methods (metaSPAdes) and will be less common once long-read sequencing of metagenomes becomes commonplace.

### Functional coverage is limited by shallow sequencing

Analysing the functional potential of bacterial communities is a common reason to prefer metagenomics instead of 16S rRNA gene amplicon sequencing. Therefore, we investigated the sequencing depth required for functional analyses at the level of both pathways and genes.

First, we investigated the impact of sequencing depth on pathway completeness, measured as percentage of detected KOs involved in a pathway. Only those pathways occurring in the reference genomes of the respective mock community were considered (Supplementary **Fig. S11**). Of the 178 pathways included in the analysis by KEGG-Decoder^26^, predicted proteins for 121 (Mock-even-70, Mock-stag-70) and 118 (Mock-stag-24) pathways were found in the reference genomes. A maximum of 80 pathways were complete at 10 Gb for Mock-even-70, 77 for Mock-stag-24, and 81 for Mock-stag-70 with the two assembly methods tested (Supplementary **Fig. S11**). One pathway was not detected in the mock communities with staggered distributions despite being present in the reference genomes, and 41, 40 and 39 pathways remained incomplete in Mock-even-70, Mock-stag-24 and Mock-stag-70 respectively. The average KEGG pathway completeness increased with sequencing depth (**Fig. 3A**). Pathway completeness reached >80 % already at 2 Gb for Mock-stag-70 and at 5 Gb for the other two mock communities. It then plateaued, with only a slight increase at maximal sequencing depth (Mock-even-70, +7.79 %; Mock-stag-24, +3.69 %; Mock-stag-70, +3.95 %).

**Fig. 3:**
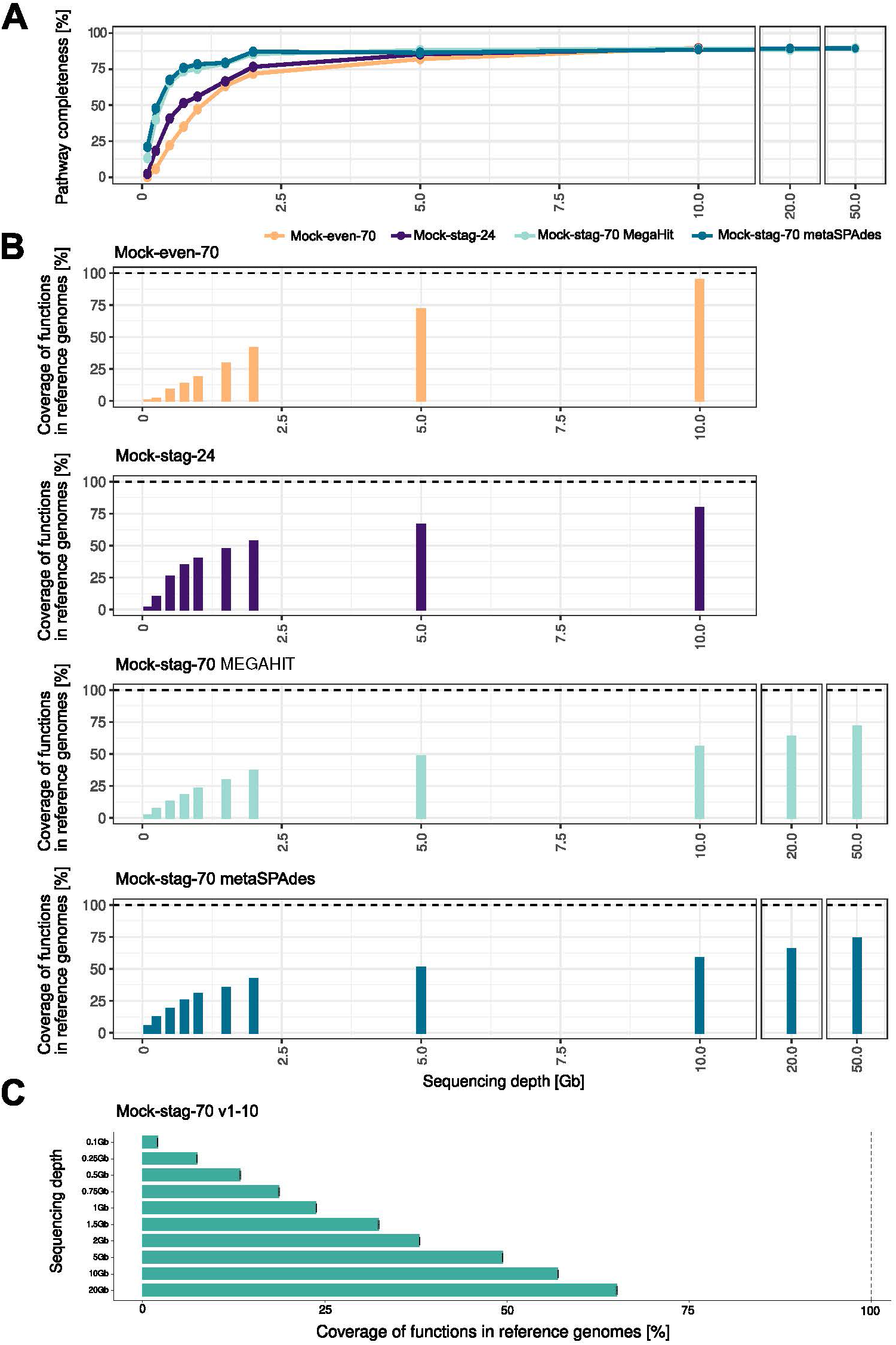
Functional coverage. **A** Average KEGG pathway completeness of Mock-even-70, Mock-stag-24, and Mock-stag-70 using contigs assembled with MEGAHIT, and Mock-stag-70 using contigs assembled with metaSPAdes. **B** Fraction of predicted proteins from the reference genomes covered by the metagenomic assemblies (MEGAHIT) for three different mock communities at the respective sequencing depths. Reads of Mock-stag-70 were additionally assembled using metaSPAdes (bottom). **C** Average coverage of functions of the reference genomes by ten *in-silico* datasets, subsampled to ten different sequencing depths each from Mock-stag-70 sequenced at 50Gb. Bars are mean values; whiskers are standard deviations. Statistics: Kruskal-Wallis rank-sum test with Benjamini-Hochberg correction (p-values in **Supplementary Table S5**).

To account for functionally unassigned proteins, the protein sequences detected at each sequencing depth were compared to the reference genomes protein repertoire (Mock-70: 227,543 protein sequences; Mock-stag-24: 82,992). The coverage of the predicted protein sequences in the reference genomes within a metagenome continued to increase up to 94.67 % for Mock-even-70 (+ 22.9 % between 5 and 10 Gb), but started to plateau at 5 Gb sequencing depth for the less diverse Mock-stag-24 (+ 12.9 % between 5 and 10 Gb) (**Fig. 3B**). In Mock-stag-70, the protein sequence coverage reached 55.5 % (MEGAHIT) and 58.3 % (metaSPAdes) at 10 Gb and increased only to 71.8 % (MEGAHIT) and 73.8 % (metaSPAdes) at highly deep sequencing (50 Gb). The protein sequence coverage obtained with the three mock communities sequenced separately was confirmed statistically using ten Mock-stag-70 subsampled bioinformatically for each sequencing depth (0.1 – 20 Gb) from the 50 Gb dataset: the recovered functions increased with sequencing step and variations between the replicates was low (**Fig. 3C**) (p-values are provided in **Supplementary Table S5**).

In summary, while a moderate sequencing depth of 5 Gb was sufficient for functional analysis at the pathway level for all mock communities, analysis at the level of single protein sequences requires greater sequencing depth depending on the diversity and distribution of the microbial community under investigation.

### Sample processing protocol and background DNA impact shallow sequencing results

To account for the influence of wet-lab factors on shallow metagenomics results, the DNA libraries were prepared in two facilities using different protocols (see methods). In addition, the two community types (Mock-even-70 and Mock-stag-24) were spiked or not with background DNA isolated from the gut content of germfree mice to simulate host DNA.

PCA plots based on relative abundance profiles showed distinct clustering due to both factors (background DNA, facility), with more pronounced effects along PC1 linked to differences in library preparation protocols between facilities (**Fig. 4A**). Within a condition (background DNA /facility pair), the extreme sequencing depths (0.1 and 10 Gb) tended to be most distant from each other. The relative abundance profiles of Mock-stag-24 prepared in facility 1, which used more template DNA (100 vs. 1 ng) and a lower number of PCR cycles (5 vs. 12), were less sensitive to the effects of sequencing depth when background DNA was present (**Fig. 4A, lower panel**).

**Fig. 4:**
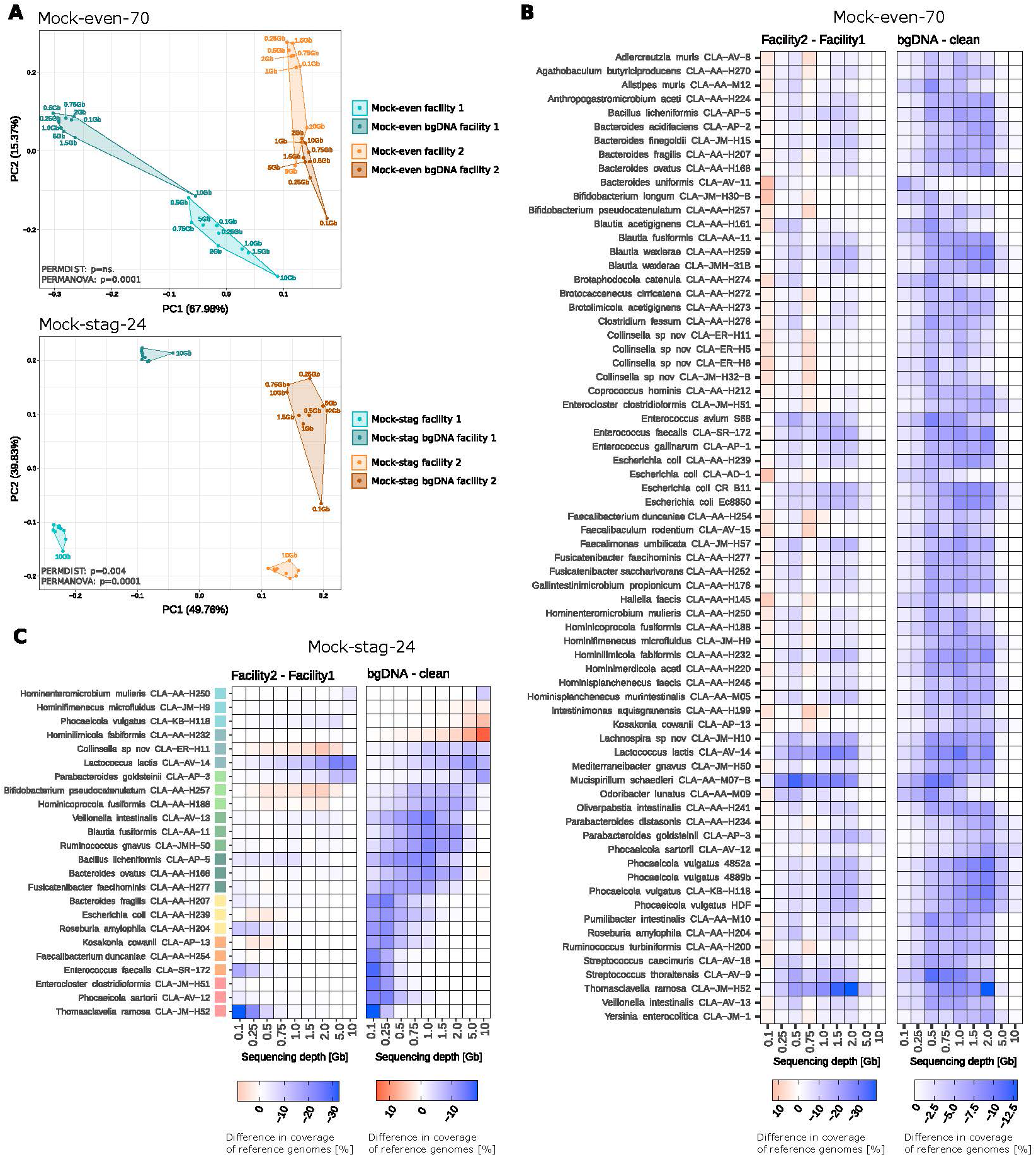
Effects of background DNA (bgDNA) and library preparation in two facilities. **A** PCA plot of relative abundance profiles in Mock-even-70 (top) and Mock-stag-24 (bottom). **B** Difference in coverage of reference genomes between facility 1 and facility 2 (left) and samples with or without bgDNA (right) for Mock-even-70. **C** Same as in panel B for Mock-stag. The reference genomes were ranked (from top to bottom) according to increasing DNA amount in the mixture, as indicated by the colour gradient (from blue to red; 0.04, 0.4, 1, 2, 4, 10, 20, 40 ng).

The effect of wet-lab factors on reference genome coverage was then assessed by calculating delta-values between the two facilities or the presence/absence of background DNA. For Mock-even-70, consistent profiles were observed across nearly all species at the 5 and 10 Gb sequencing depths, regardless of facility or background DNA (**Fig. 4B**). In contrast, variations in reference genome coverage of up to 39 % for facilities, and up to 13 % for background DNA were observed at sequencing depths ≤2 Gb. For Mock-stag-24, the variation between samples from the two facilities was additionally influenced by the initial DNA amount indicated as coloured boxes in **Fig. 4C**. With a reference genome input >1 ng and a sequencing depth >1 Gb, the total variance in reference genome coverage between the two facilities dropped below 3.69%. A similar trend was observed for the impact of background DNA on the coverage of reference genomes in Mock-stag-24, but with a more pronounced influence of the amount of DNA. The effect of background DNA was negligible (total variance 1.33 %) with ≥10 ng template DNA and a sequencing depth ≥1.5 Gb. For a DNA amount >1 ng per strain in Mock-stag, a sequencing depth of 5 Gb or higher was necessary (total variance 3.23 %).

Analysed per strain, the relative abundance profiles in Mock-even-70 were consistent between the two library preparation strategies, except for marked differences for *Collinsella* sp. CLA-ER-H5 due to sequencing depth in facility 1 (**Supplementary Fig. S5A**). This indicates that the differences shown in Figure 4A are due to changes in the relative abundance of a few strains. For Mock-stag-24, deviation from the theoretical relative abundance values was different between the two facilities (**Supplementary Fig. S5B**). The relative abundance of *E. clostridioformis*, for example, matched the expected relative abundance in facility 1, while it had the highest relative abundance in facility 2. The most abundant species in facility 1 was *T. ramosa*. This indicates that library preparation (specifically the amount of template DNA and number of PCR cycles) influenced relative abundance profiles. (**Supplementary Fig. S5B**)

We further analysed the effects of the wet-lab factors on the predicted protein profiles. Overall, facility 1 provided a more complete functional profile for both mock communities (**Supplementary Fig. S12**). The addition of background DNA was associated with a decrease in coverage, which was attenuated at high sequencing depths in both mock communities prepared in facility 1. In general, the influence of background DNA was more pronounced with the protocol used in facility 2.

In summary, library preparation with a higher amount of template DNA and fewer PCR cycles is recommended for more robust taxonomic and functional profiles. If necessary, a higher sequencing depth can compensate for the effects of lower DNA amount or the presence of non-target DNA.

## Discussion

In this work we investigated the potential and limitations of shallow metagenomic sequencing using complex mock communities and sequencing depths from 0.1 Gb (0.3 million reads) to 50 Gb (166.7 million reads), with a specific focus on strain-level and functional analysis.

Previous studies reported that taxonomic profiles based on shallow shotgun sequencing (0.5-4 million reads) can reflect those obtained at a higher sequencing depths^2,3,6,8^. Most of those studies relied on deep sequencing of native samples then subsampled *in-silico*. Using defined input, we have confirmed these results, showing that shallow metagenomics is suitable for reference-based taxonomic analysis: reads for all strains were detected already at 0.1 Gb (0.3 million reads), even for lowly abundant taxa in the staggered communities, enabling prevalence and relative abundance analyses despite overall low genome coverage. This is in line with the study of Hillmann et al. and Xu et al. who reported that 0.5 million sequences (= 0.15 Gb) and 0.5 Gb, respectively, was sufficient for taxonomic analysis of the human gut microbiome^2,3^. As most of the metagenomic costs is due to sequencing, being able to pool more samples on the same run reduces the expenses, making shallow metagenomics at 0.5 Gb/sample attractive for diversity and composition profiling in studies with large number of samples. However, accurate analysis necessitates a high-quality, reference database of individual genomes.

Metagenomics is useful for high resolution taxonomic analysis, which is limited with 16S rRNA gene amplicon sequencing. In a previous study using reference-based analysis, species profiles at 0.5 million reads (0.15 Gb) showed an average correlation of 0.99 with ultra-deep sequencing (750 Gb)^3^. Using mock communities, Xu et al. recovered 62 % of the 62 species at 1 Gb/sample using MetaPhlAn2 and the Refseq database^2^. The analysis of our mock communities with MetaPhlAn4 at 1 Gb showed similar result (Mock-stag-24: 75%, Mock-even-70: 67%, Mock-stag-70: 43%), without resolution at the strain level (four strains of each *E. coli* and *P. vulgatus* in Mock-even-70). Using the reference genomes, we detected reads for all 70 and 24 strains in the mock communities at the lowest sequencing depth of 0.1 Gb. The reference-based detection and discrimination of different strains with an ANI of up to 99.66 % was possible in our study. However, sequencing depths >5 Gb are required to obtain nearly full genome coverage, which must be adjusted depending on the expected abundance and genome size of the targeted strains in complex samples.

A sequencing depth ≥5 Gb was required to reconstruct sufficient MAGs, which agrees with previous findings^3,6^. Importantly, at high sequencing depth (10 to 50 Gb), more MAGs were created than input strains, yet not all strains were represented by a MAG. Inspecting this further revealed a substantial number of hqMAGs that were chimeric due to containing fragments assigned to more than one reference genome. This occurred despite excluding contigs smaller than 1,500 bp during binning to prevent false inflation of chimerism. These results echo previous reports that ∼5 % of genes within MAGs differ from the dominant reconstructed taxa^48^. A recent study, which evaluated the quality of short-read assemblies from a natural soil community by mapping them to the respective long-read data, reported that assembly failures occurred in most genome bins^49^, which agrees with our findings.

Our results show that deeper sequencing does not guarantee more complete MAG reconstruction and does not eliminate chimerism. Data from the staggered Mock communities indicate that having more reads for a reference genome does not assure its coverage by a MAG and does not prevent the risk of multiple alignments. Instead, we observed that the creation of several MAGs for the same reference genome was common. The aforementioned study of a natural soil community also found that overabundance of reads can lead to high-coverage misassemblies^49^. Bioinformatic approaches that reduced the occurrence of chimeric hqMAGs in our work were multi-coverage binning and strain aware assembly, which is in line with previous studies^34,49^. Long-read sequencing also increased the number of coherent hqMAGs. This highlights the need to consider long-read or hybrid sequencing for projects where the goal is strain reconstruction, whilst simultaneously recognising the need to also enhance such approaches further^50^. Nevertheless, none of the approaches tested could resolve the issue completely and the data presented raises concerns about the prevalence of spurious sequences in ever-growing MAG catalogues^51^, as we used common bioinformatic workflows to create them.

To quantify the functional information lost with decreasing sequencing depth, we calculated the average KEGG pathway completeness, showing that >2 Gb is required to reach >50 % completeness. In contrast, one study with a mock community of 62 human gut bacteria reported that the functional description of shallow metagenomic samples (1 Gb) was “quite similar” to the one obtained from the reference genomes at KEGG level 1 and 2^2^. Another study based on in-silico subsampling of an ultra-deep metagenome (2.5 billion reads; 750 Gb) to 0.5 million reads (0.15 Gb) showed a nearly full recovery of KEGG Orthology groups^3^. This result might be influenced by differences in the complexity of analysed communities and by the subsampling process being rarefied repeatedly and therefore creating a higher chance of recovering functions. At the level of predicted protein sequences, our data suggests that shallow metagenomics is not advisable, as a recovery of <75 % predicted proteins was reached at sequencing depths ≤5 Gb. This agrees with a previous study of antimicrobial resistance genes at different sequencing depths, in which at least 80 million reads (24 Gb) was needed to recover the full richness of AMR gene families^52^.

Regarding the influence of wet-lab factors, the performance of library preparation procedures has previously been compared for traditional metagenomic sequencing of native samples and Mock communities, revealing that the community diversity, amount of input DNA, and the sequencing platform alter the results^53–55^. When analysing the effects of library preparation in two facilities and the addition of background DNA from the gut content of germfree mice, both factors impacted taxonomic and functional results at the sequencing depths <5 Gb. Strains with a higher relative abundance were less sensitive to these effects, even at lower sequencing depths. If the use of different protocols in large studies cannot be avoided (e.g., when handling low biomass samples), or a high amount of background DNA is to be expected, higher sequencing depth may improve the robustness of results.

This study has some limitations: (i) Metagenomics can be used to study various environments, which may require different strategies; due to our own research interests, we focused on wet-lab parameters related to gut microbiomes. Although we could show their overall effects on metagenomic data, the study design was not appropriate to disentangle the effect of each parameter separately. Previous studies have investigated other aspects such as low biomass, contamination, or additional library preparation protocols (DNA input, PCR cycles, fragment/insert size)^53,56^; (ii) Whilst we used MetaPhlAn4 for some of the analyses, most results were obtained using the exact strains (*i.e.*, their genomes) as a reference, which represents an ideal case for taxonomic and functional readouts; (iii) We have tested multiple approaches, including standard workflows used by many, to study the chimeric MAGs. We do not exclude that additional strategies not tested in this work may perform better. Future benchmarking of bioinformatic tools, as in Critical Assessment of Metagenome Interpretation (CAMI), will be helpful to move forward with comprehensive testing of new approaches to address chimeric MAGs at scales that go beyond our single study^57^; (iv) the mock communities analysed included only bacteria; whilst bacteria are dominant members in many microbiomes, the work excludes the influence of other types of microbes; (v) We used a higher number and more complex mock communities (up to 70 strains, including different distributions) than in previous studies; they nonetheless are only approximations of native bacterial populations in gut microbiomes, which include ca. 300 species, with 150-200 usually detected by sequencing.

In summary, due to the lack of studies benchmarking shallow metagenomics using complex mock communities, potential users may not be aware of the pitfalls of this approach. The sequencing depth needs to match the main aims of a study. Factors to be considered include the expected microbial diversity in the target samples, the evenness of distribution within the communities, and the amount of DNA available. Shallow metagenomics is a valid choice for a large study aiming at database-guided taxonomic profiles of well-characterised environments. However, it is not appropriate for high-resolution functional analysis and *de-novo* strain resolution. Even at high depths of sequencing, MAG-based approaches require careful planning and execution to minimise the generation and spread of artificial bacterial diversity.

## Acknowledgement

We are grateful to: (i) Evelyn Deis for sample processing, Alina Viehof-Beckmann for providing gut content of germ-free mice, and Ntana Kousetzi for outstanding support with molecular work (all from Functional Microbiome Research Group, University Hospital of RWTH Aachen); (ii) Christel Driessen for genomic library preparation (Department of Medical Microbiology, Infectious Diseases and Infection Prevention, Maastricht University Medical Centre). This work was supported by: (i) the Genomics Facility of the Interdisciplinary Center for Clinical Research (IZKF) Aachen within the Faculty of Medicine at RWTH Aachen University; (ii) the DFG-funded NGS Competence Center Tübingen (INST 37/1049-1) and the Institute for Medical Microbiology and Hygiene at the University Hospital (Tübingen, Germany), including help by the Quantitative Biology Center (QBiC) for raw data management and storage; (iii) the Life Science Computer Cluster (LiSC) of the University of Vienna for the processing of long-read sequencing data.

## Authors contributions

NT: conceptualization, methods and software, investigation, formal analysis, validation, interpretation, visualization, data curation, writing – original draft; CP: methods and software, validation, interpretation, writing – review and editing; JS and PP: investigation, formal analysis, writing – review and editing; DB and JP: resources, funding acquisition, writing – review and editing; TCAH: conceptualization, interpretation, writing – original draft; TC: conceptualization, visualization, interpretation, resources, supervision, project administration, funding acquisition, writing – original draft.

## Material and data availability

The raw metagenomic sequencing data was deposited at the European Nucleotide Archive/NCBI and is accessible under Project no. PRJEB83573.

The code used for bioinformatic analyses is available on GitHub: https://github.com/ClavelLab/Benchmarking-shallow-Metagenomics

All strains (and their DNA) used in this study are available at the Leibniz Institute DSMZ (German Collection of Microorganisms and Cell Cultures): www.dsmz.de/miBC; https://hibc.rwth-aachen.de/.

## Ethics

All mice experiments were performed under Ethical Approval (LANUV no. 81-02.04.2023.A253) in accordance with the German Animal Protection Law (TierSchG).

## Online Supplementary Files

**Supplementary Fig. S1:**
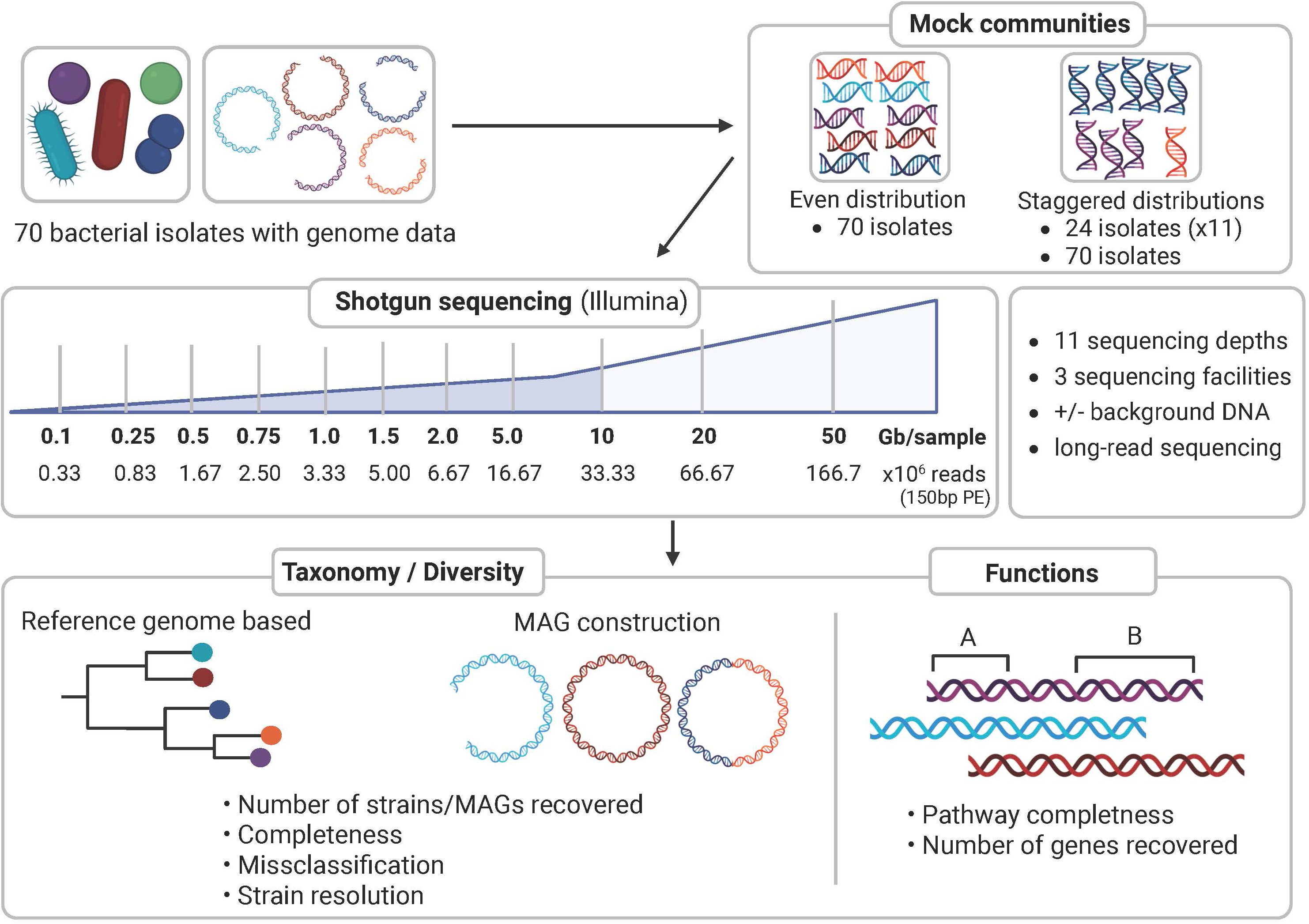
Schematic overview of the experimental design. The genomic DNA of 70 bacterial isolates (www.dsmz.de/miBC; https://www.hibc.rwth-aachen.de) was used as input to create 13 different Mock communities. Two Mock communities contained all members, one each with an even distribution (Mock-even-70) or staggered distribution (Mock-stag-70). Additionally, mock communities with varying staggered distribution of 24 isolates were created: Mock-stag 24, sequenced at nine different sequencing depths; Mock-stag 24 v1-10 sequenced at 10 Gb for testing multi-coverage binning for MAG construction. A second version of Mock-even-70 and one Mock-stag-24 were created by spiking DNA isolated from the gut content of germfree mice. Libraries for these four mock communities (Mock-even-70 and Mock-stag-24 ± bg) were prepared in two different sequencing facilities. A library per sequencing depth was then sequenced using the Illumina technology (short reads) at up to 11 sequencing depths. Mock-stag-70 was also sequenced using Oxford Nanopore Sequencing (long reads) in a third facility. Bioinformatic analyses included: (i) number and relative abundance of strains; (ii) coverage of reference genomes; (iii) number and diversity of predicted proteins and completeness of functional pathways; (iv) strain-level resolution using both a reference-based and metagenome-assembled genome (MAG) approach.

**Supplementary Fig. S2:**
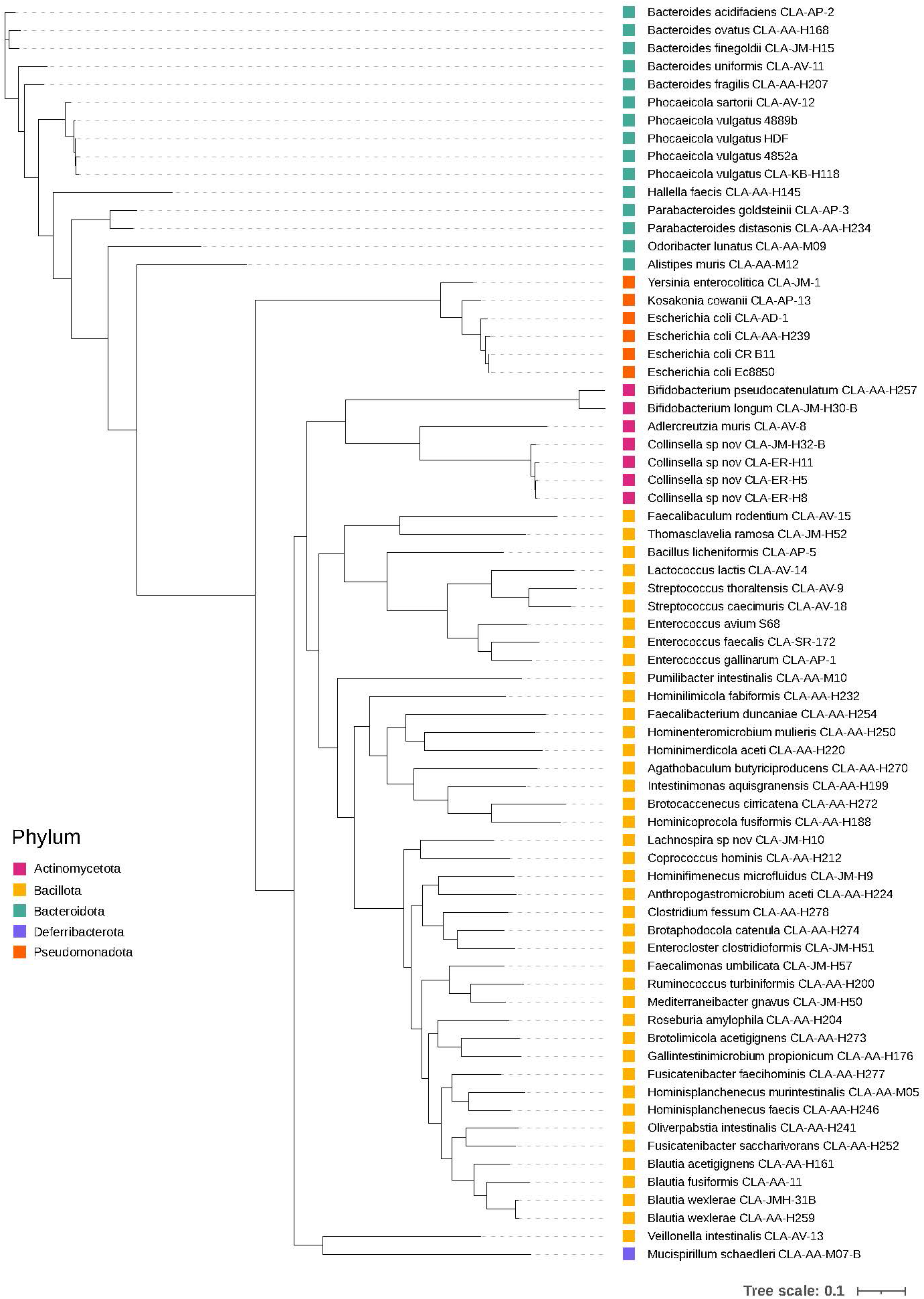
Phylogenetic tree of bacterial strains in the mock communities. A phylogenetic tree of high-quality, draft genomes of the 70 isolates was created using PhyloPhlAn v3.0.67. The phyla to which the strains belong are represented by coloured boxes at the end of the branches. Distribution of the strains in each mock community is provided in **Supplementary Table S1**.

**Supplementary Fig. S3:**
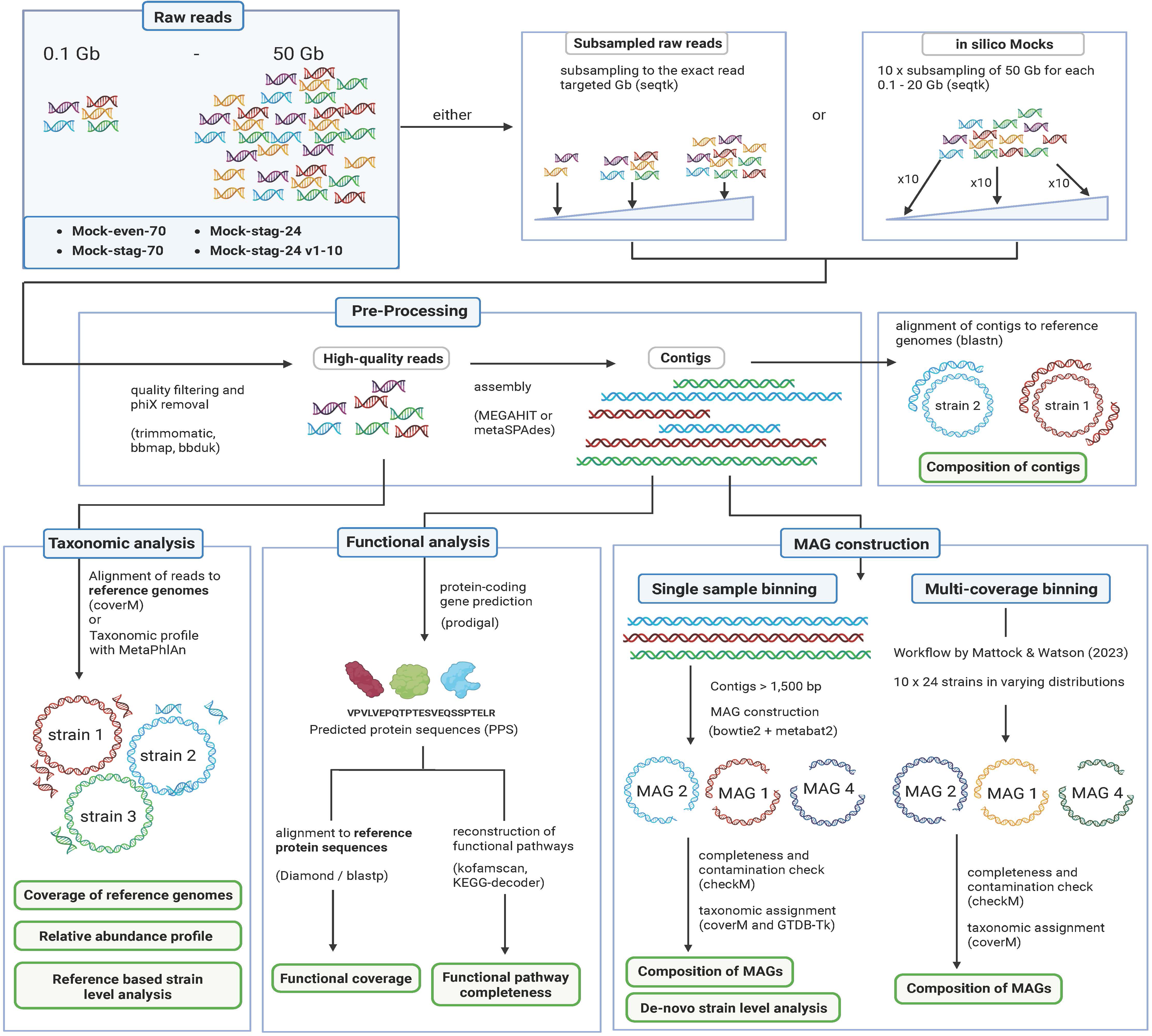
Schematics of the bioinformatics workflow. The details of each analysis are described in the methods.

**Supplementary Fig. S4:**
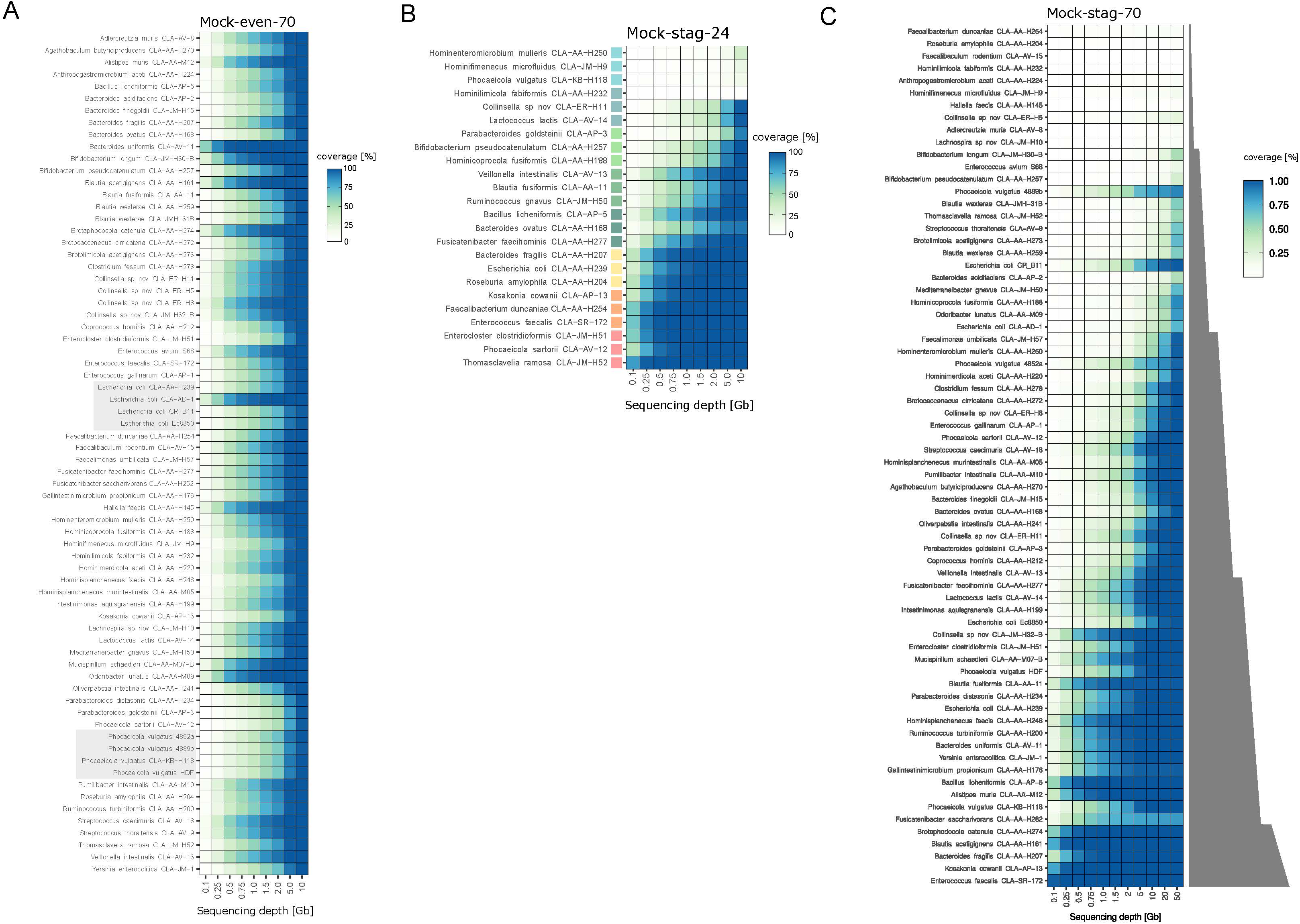
Coverage of individual reference genomes. Heatmaps showing the coverage of all reference genomes in Mock-even-70 (**A**), Mock-stag-24 (**B**), and Mock-stag-70 (**C**) by metagenomic reads at up to 11 sequencing depths (x-axis). The grey boxes in panel **A** indicate the multiple *E. coli* and *P. vulgatus* strains. The reference genomes in the mock communities with staggered distribution in panel **B** and **C** were ranked (from top to bottom) according to increasing DNA amount in the mixture, as indicated by the colour gradient in panel B (from blue to red; 0.04, 0.4, 1, 2, 4, 10, 20, 40 ng) or the grey gradient in panel C (concentrations are provided in **Supplementary Table S1**).

**Supplementary Fig. S5:**
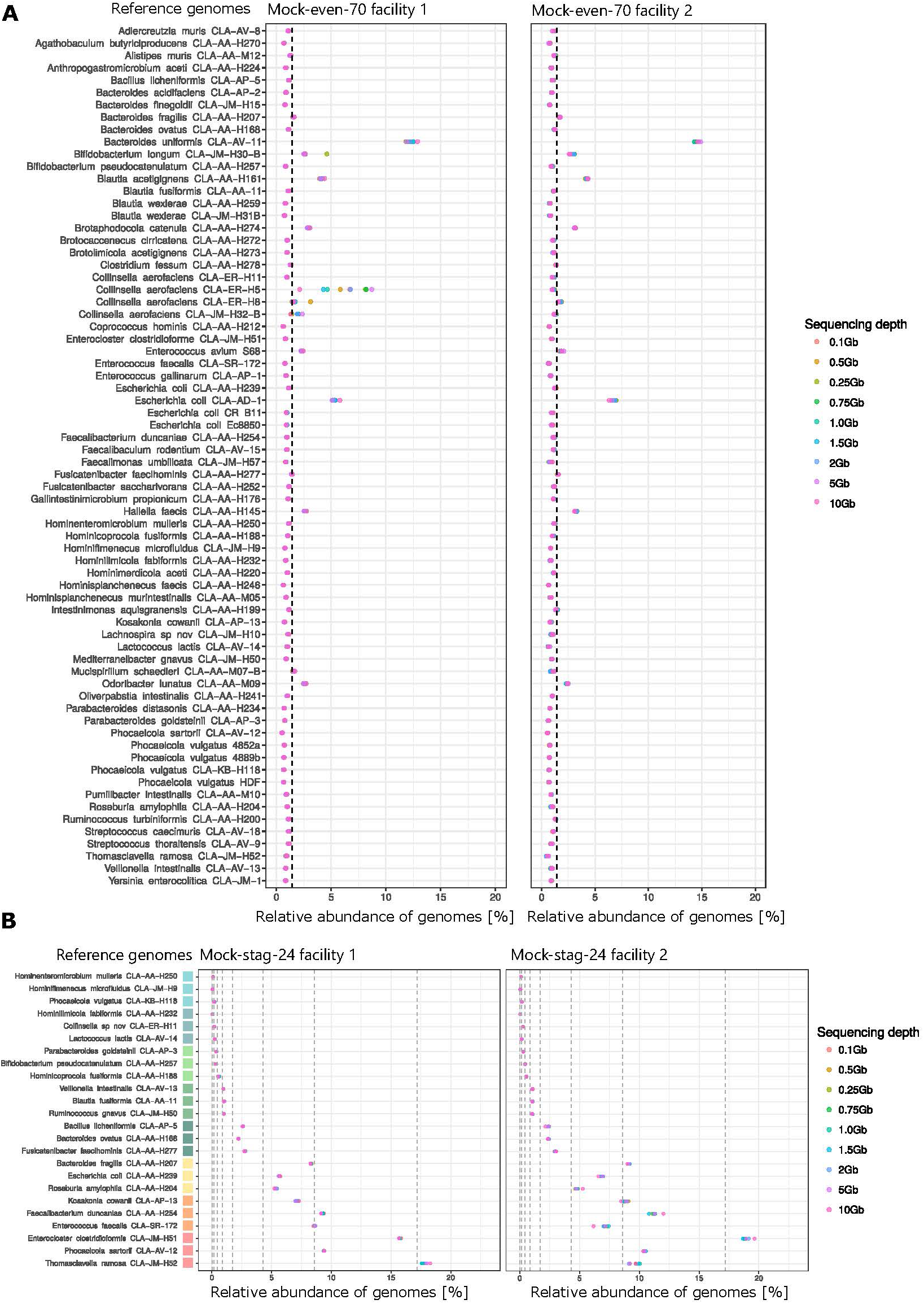
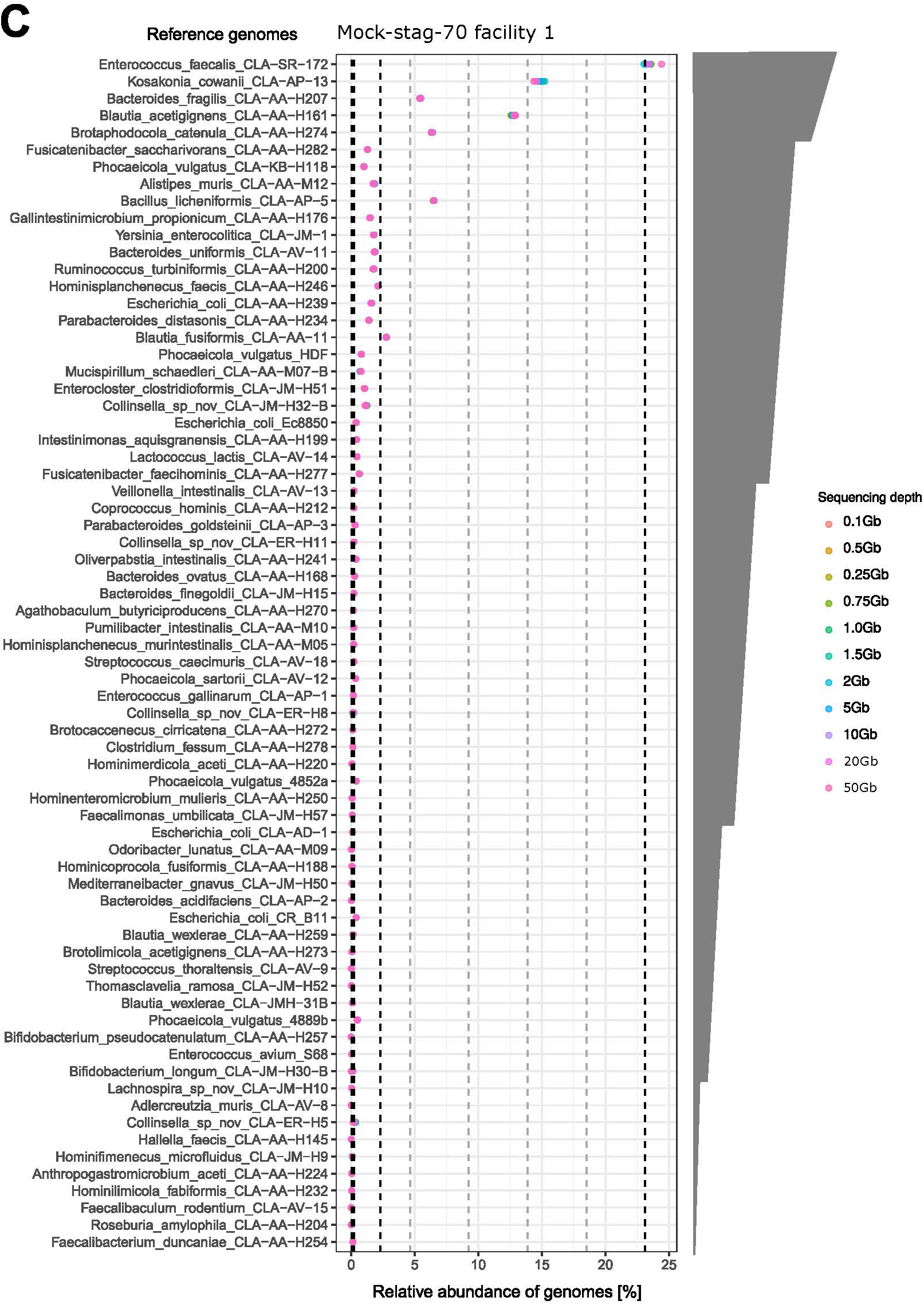
Relative abundance of reads per reference genome. Relative abundances of each strain in (**A**) Mock-even-70 in facility 1 and 2, (**B**) Mock-stag-24 in facility 1 and 2, and (**C**) Mock-stag-70 in facility 1. Metagenomic reads were assigned to the reference genomes using coverM. Sequencing depths are indicated with the coloured dots. The theoretical relative abundances are indicated by vertical dashed lines. For Mock-stag-24 in panel B and Mock-stag-70 in panel C, the reference genomes are ranked from top to bottom by increasing DNA amount in the mixture either indicated by coloured boxes (from blue to red; 0.04, 0.4, 1, 2, 4, 10, 20, 40 ng) or the grey gradient in panel C (concentrations are provided in **Supplementary Table S1**).

**Supplementary Fig. S6:**
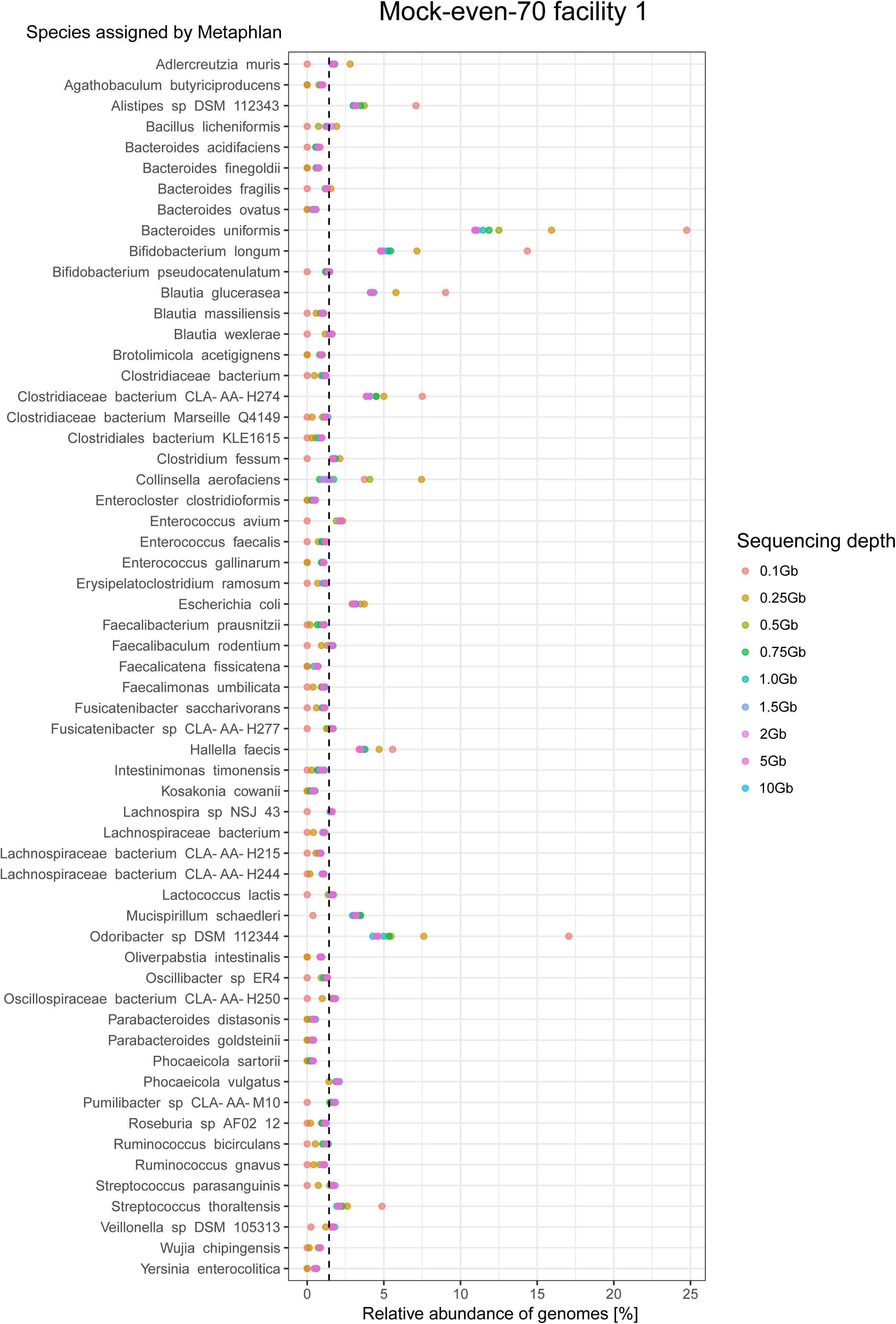
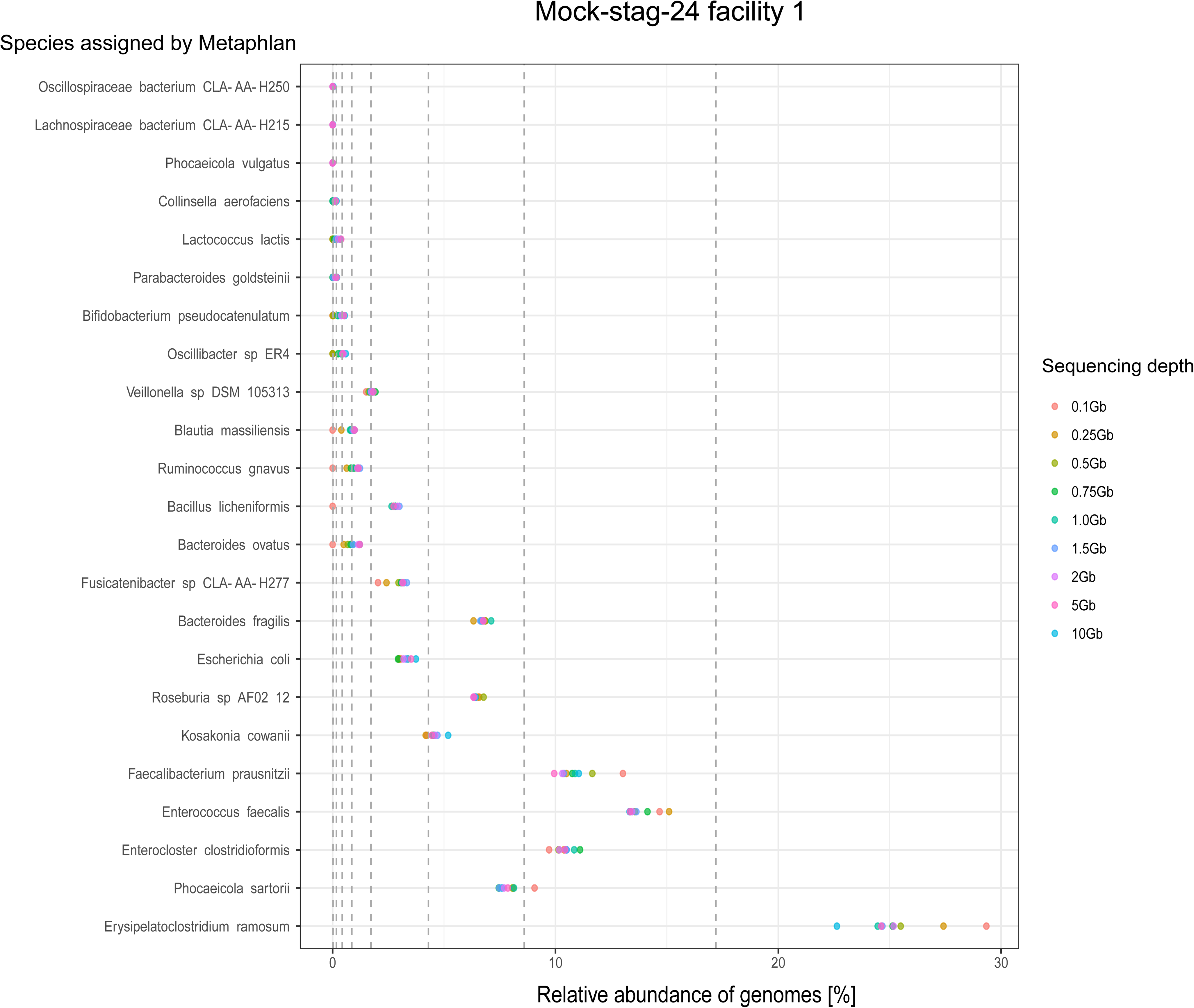
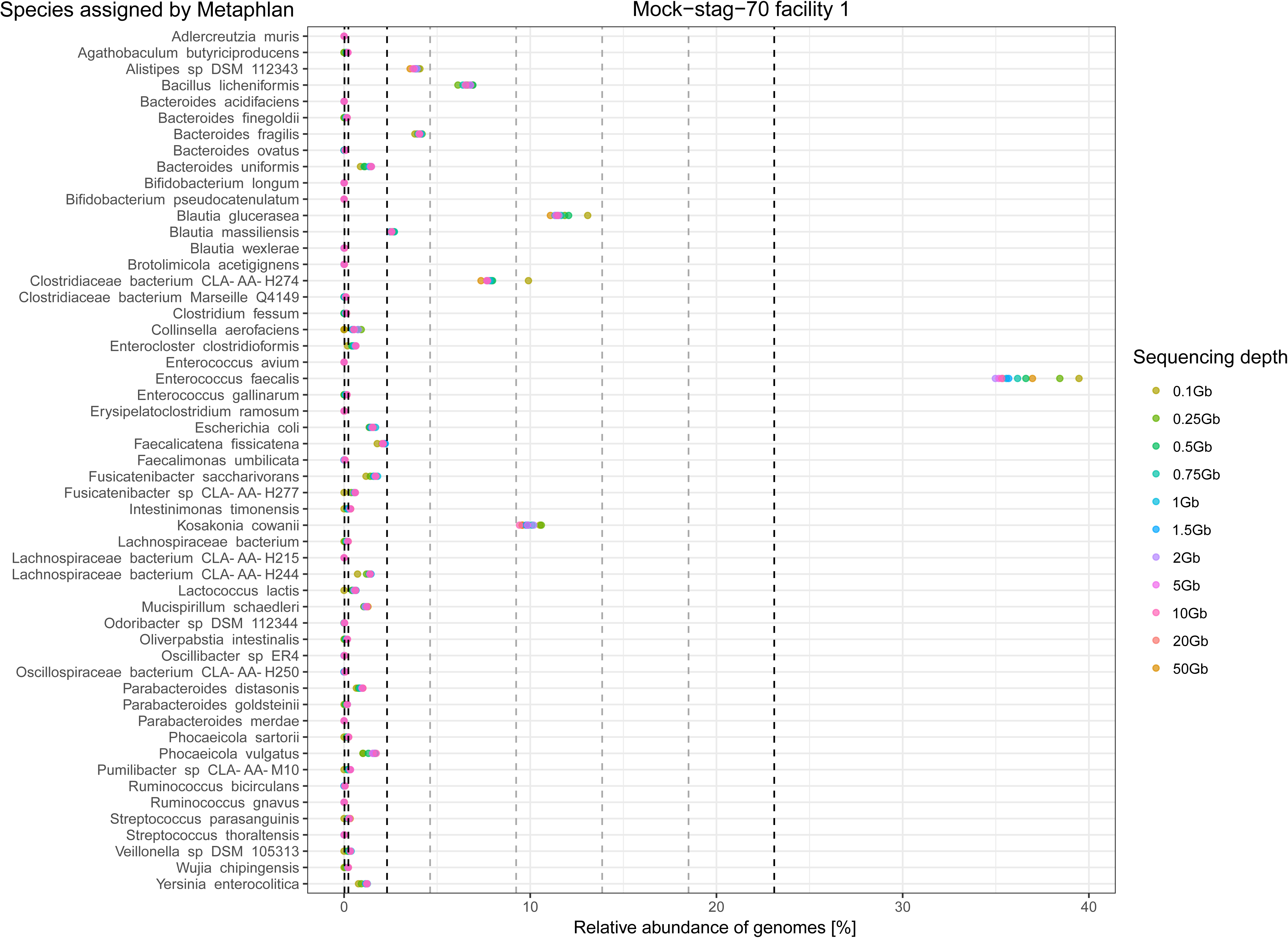
Relative abundance after “non-supervised” taxonomic assignment using MetaPhlAn4. Relative abundances of each strain in (**A**) Mock-even-70, (**B**) Mock-stag-24, and (**C**) Mock-stag-70. Metagenomic reads were assigned using MetaPhlAn4. Sequencing depths are indicated with the coloured dots. The theoretical relative abundances are indicated by vertical dashed lines. For Mock-stag-24 in panel B and Mock-stag-70 in panel C, the reference genomes are ranked from top to bottom by increasing DNA amount in the mixture (see concentrations in **Supplementary Table S1**).

**Supplementary Fig. S7:**
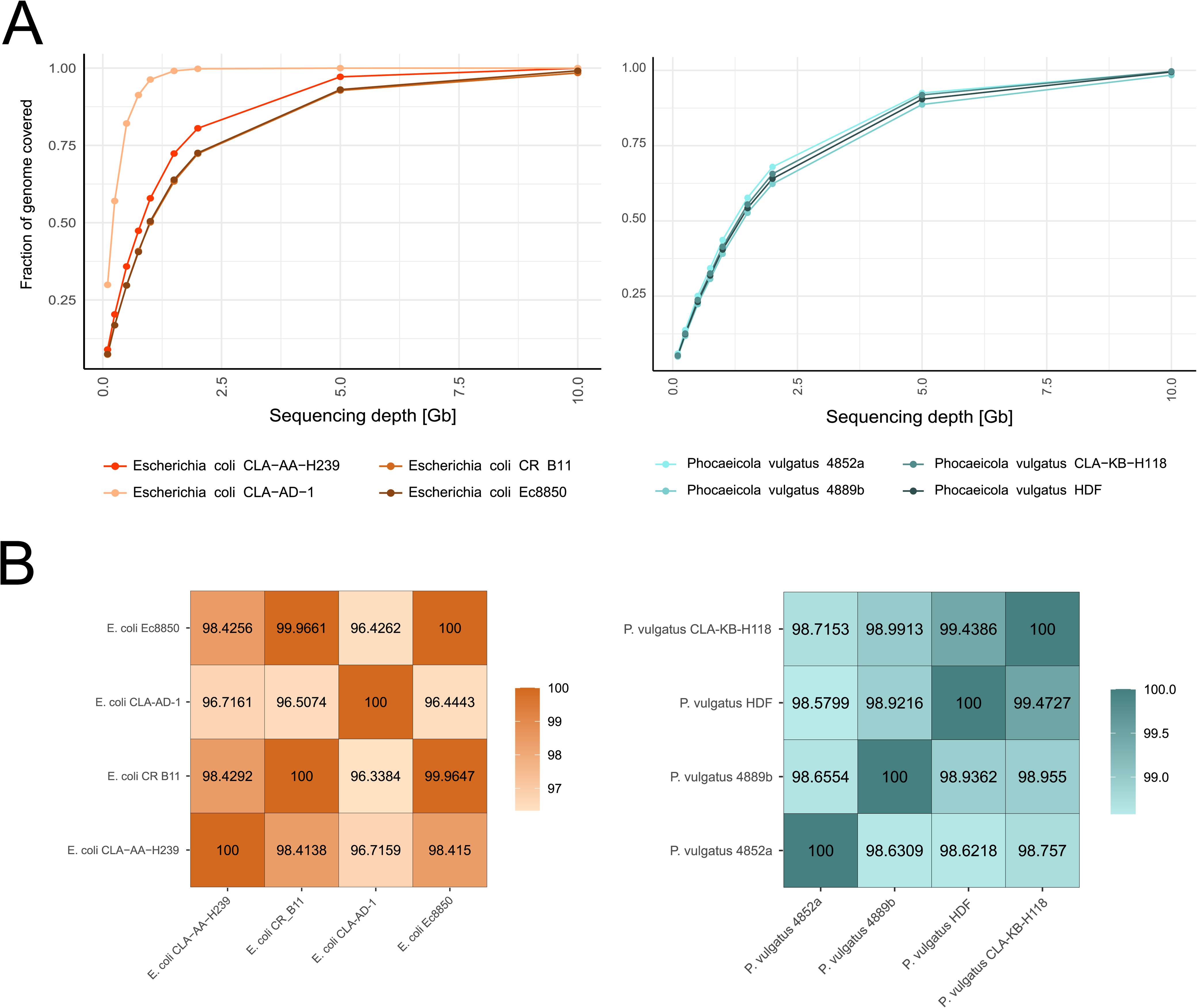
Strain level resolution based on reference genomes. **A** Reference genome coverage (y-axis) for the four strains of *E. coli* and *P. vulgatus* at the nine different sequencing depths (x-axis). **B** ANI values between the strains within each species.

**Supplementary Fig. S8:**
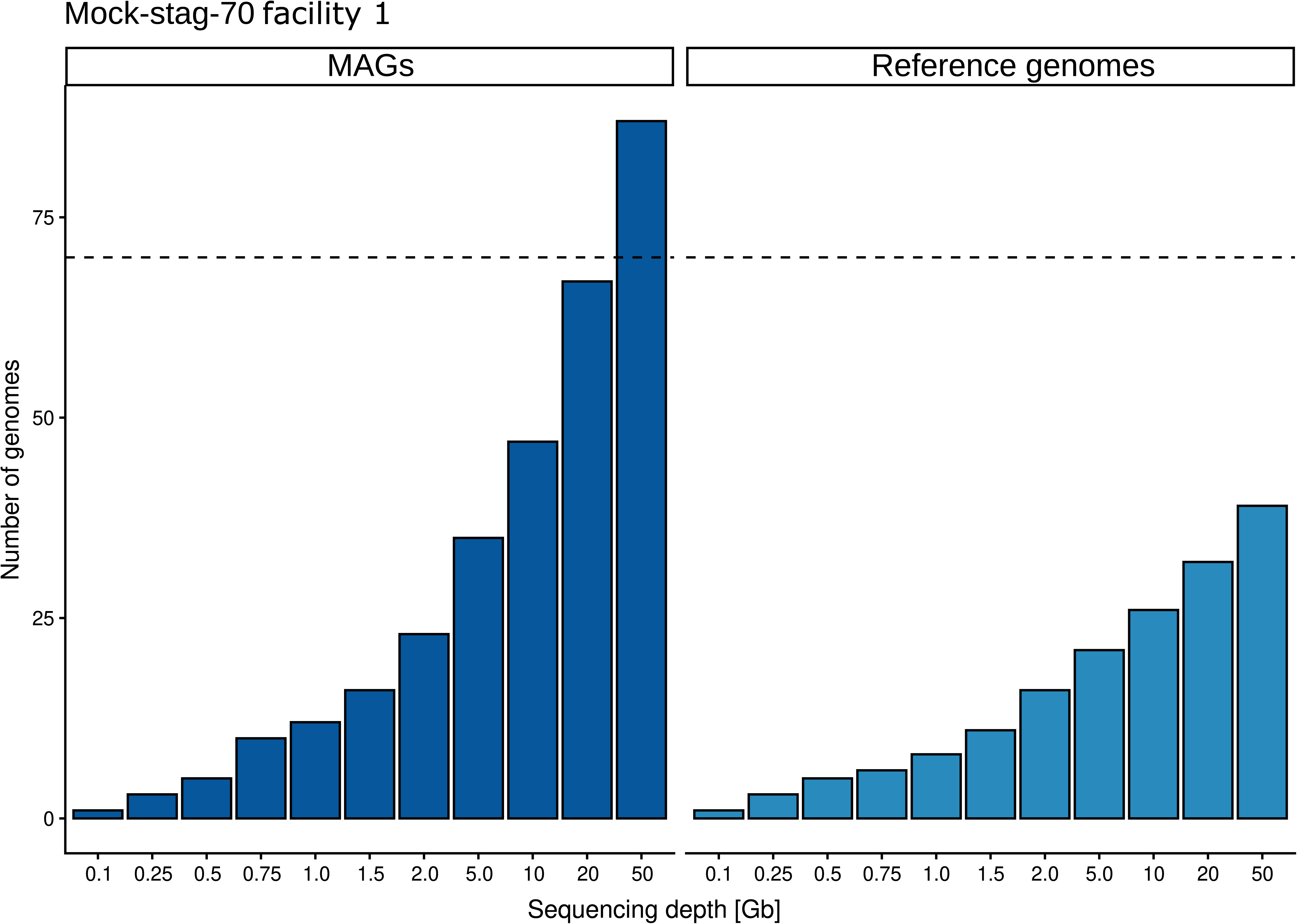
MAGs numbers in Mock-stag-70. Strain analysis based on the MAGs assembled in Mock-stag-70 using MEGAHIT for assembly at each sequencing depth individually. The left panel (dark blue) depict the number of MAGs assembled from the shotgun data; the right panel (lighter blue) show the number of reference genomes matching MAGs (coverM), indicating that multiple MAGs were reconstructed for some Mock species. The reference genome with the highest coverage by MAG reads was chosen as representative of that MAG.

**Supplementary Fig. S9:**
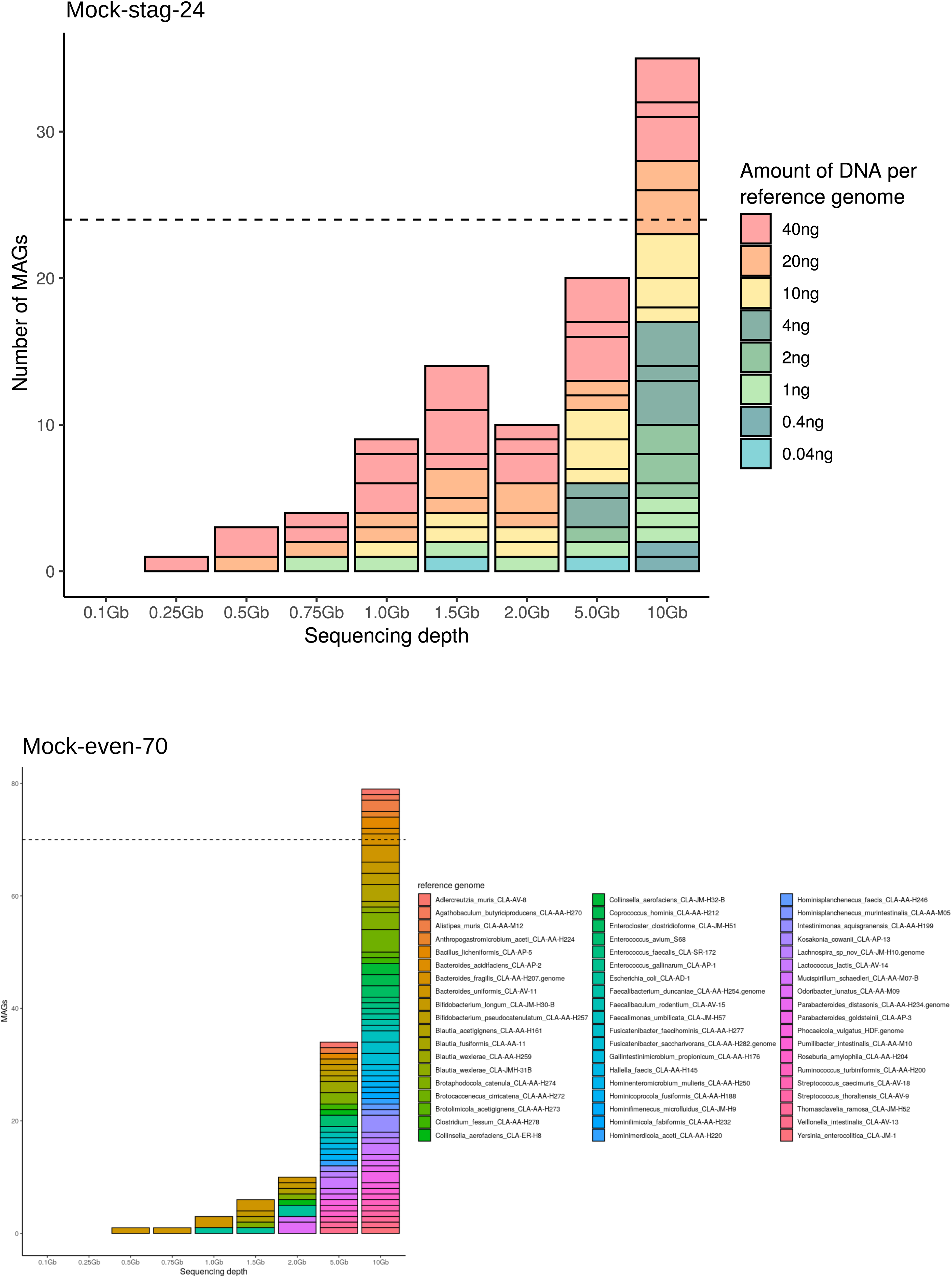
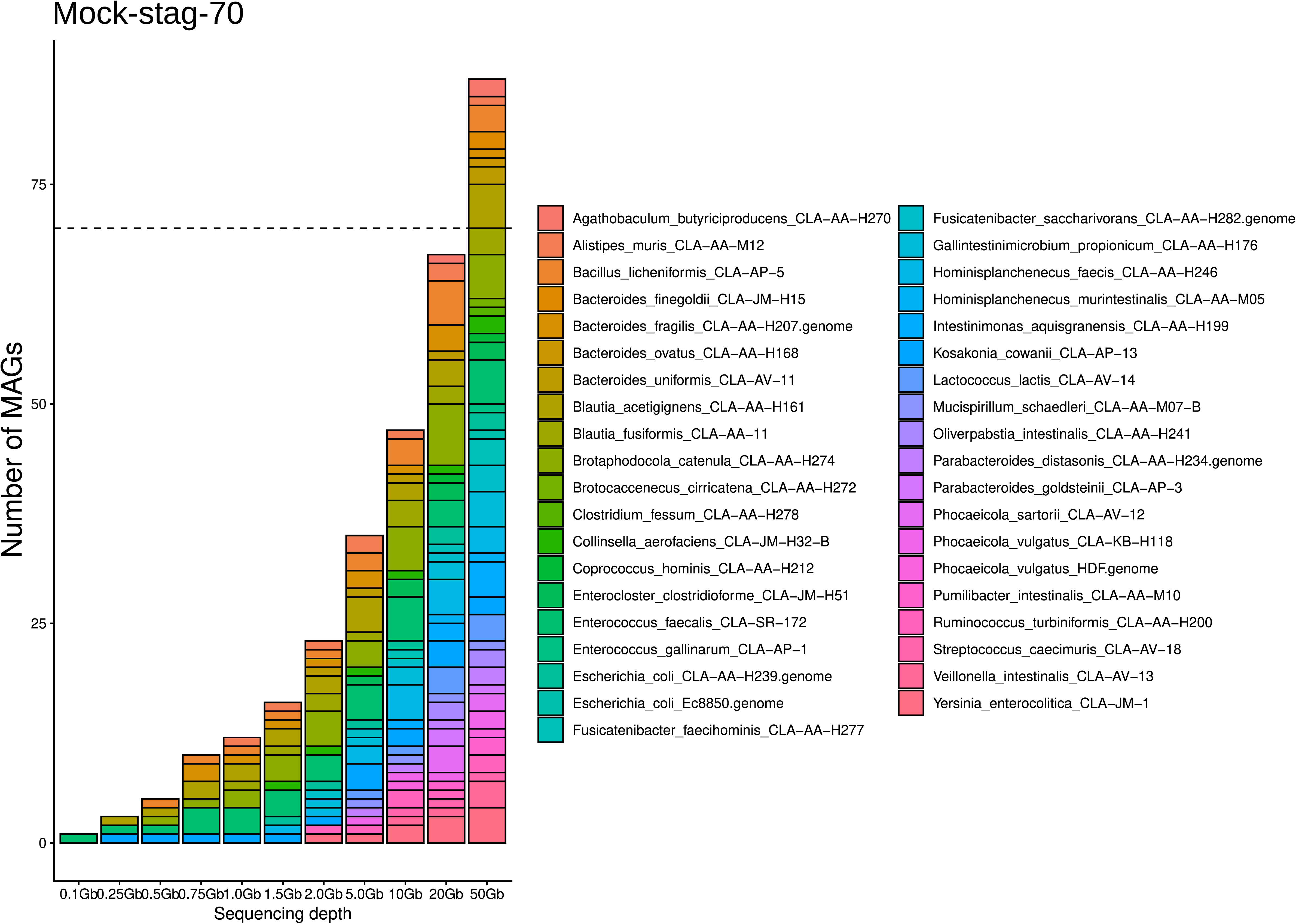
Reference genomes per MAG for Mock-stag-24, Mock-even-70, and Mock-stag-70. Stacked bar charts of the number of MAGs per sequencing depth. Each box represents one reference genome. The size of the box indicates the number of MAGs that were assigned to this reference genome. The colours in the graph for Mock-stag-24 indicate the amount of input DNA for a given strain in the pool. The colour of the graph for Mock-even-70 and Mock-stag-70 indicates the reference genomes.

**Supplementary Fig. S10:**
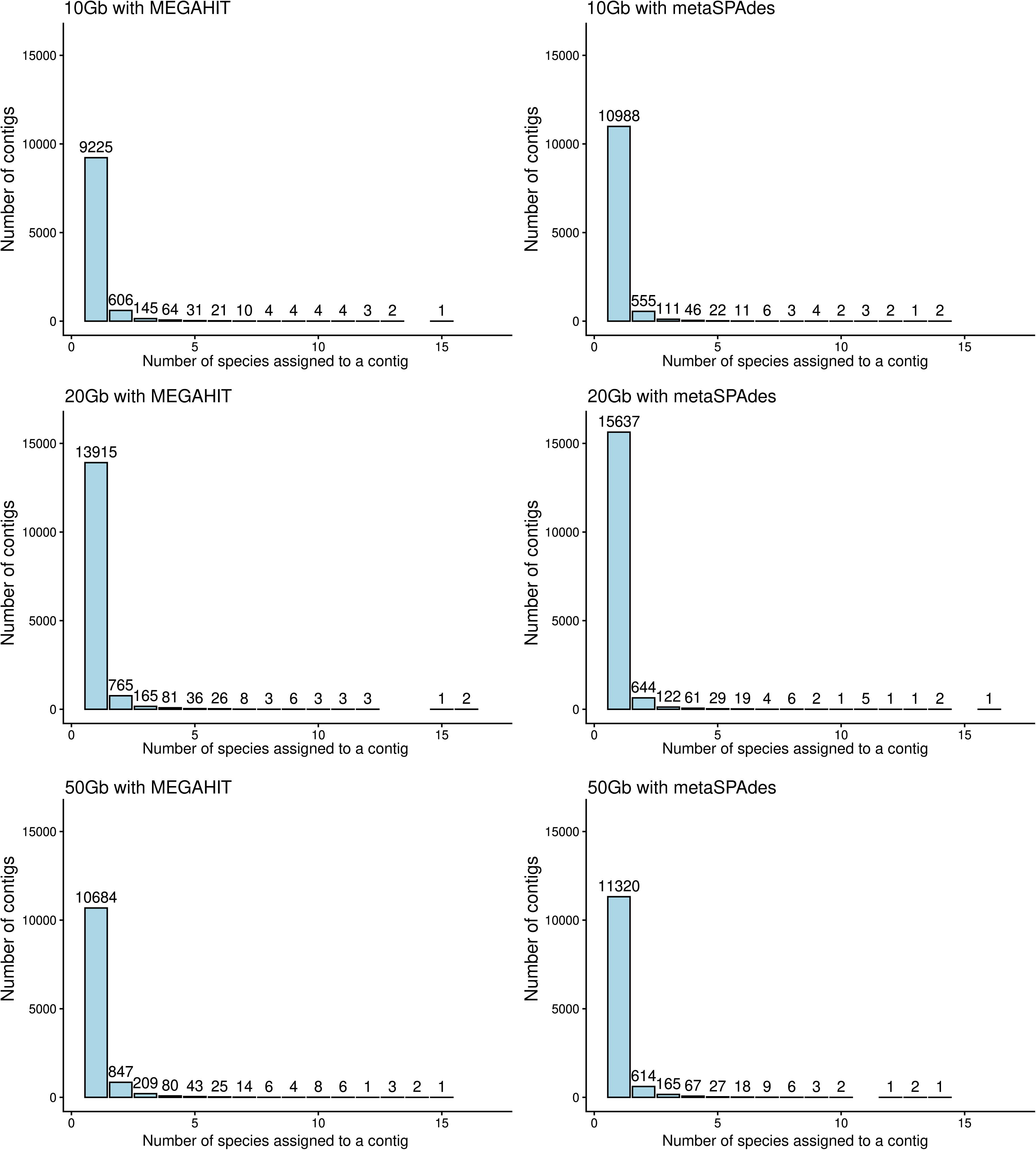
Contigs assigned to reference genomes. Contigs from metagenomes sequenced at 10 Gb, 20 Gb and 50 Gb either assembled with MEGAHIT or metaSPAdes were aligned to the reference genomes with blastn (-perc_identity 97% -evalue 1e-10, alignment length >150 bp). The number of contigs (y-axis) is plotted against the number of reference genomes assigned to a contig (x-axis).

**Supplementary Fig. S11:**
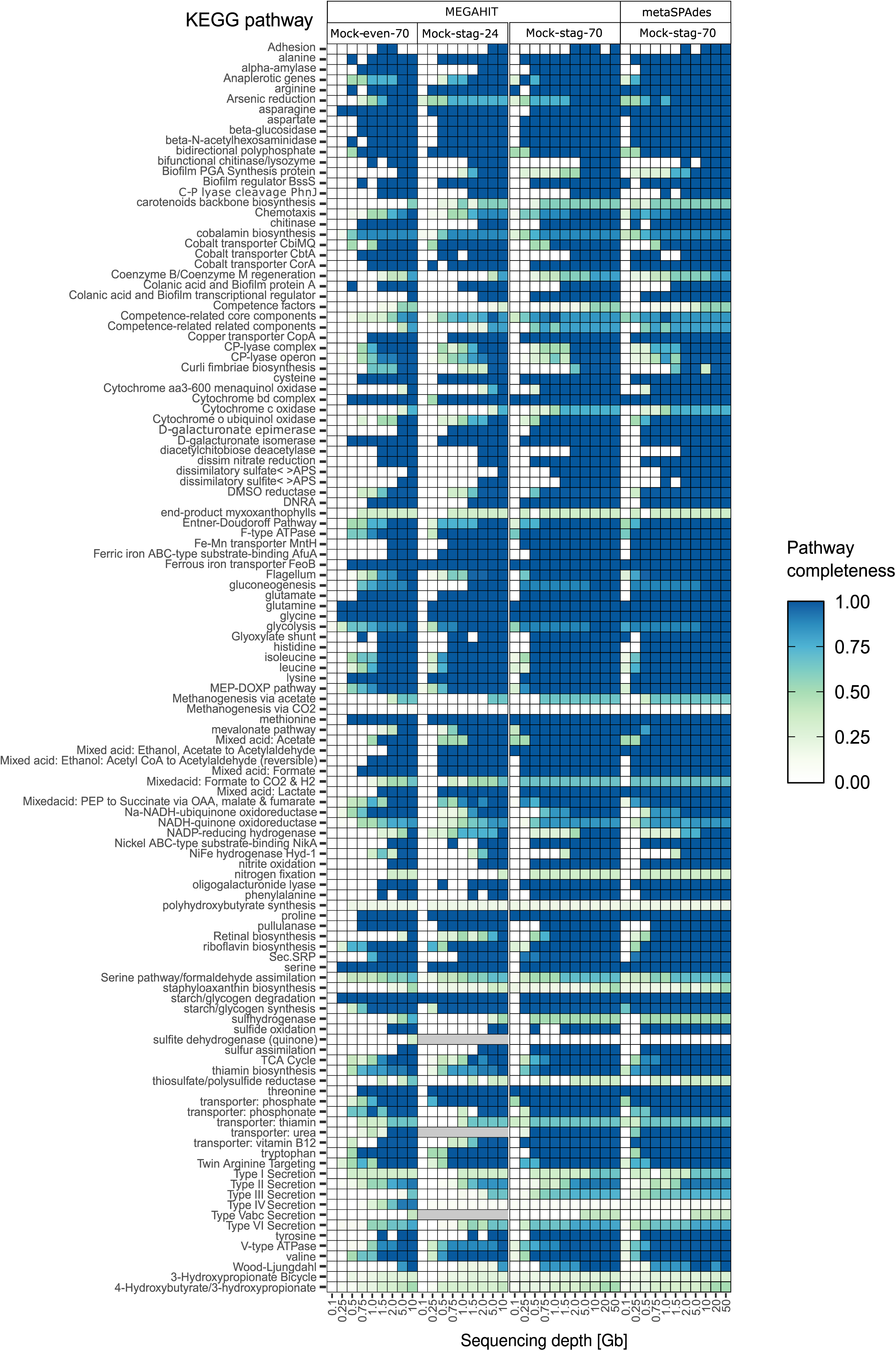
Heatmaps of functional coverage. Completeness of all KEGG pathways present in the reference genomes in Mock-even-70, Mock-stag-24 and Mock-stag-70 assembled with MEGAHIT, and Mock-stag-70 assembled with metaSPAdes (from left to right) at the up to 11 sequencing depths (columns).

**Supplementary Fig. S12:**
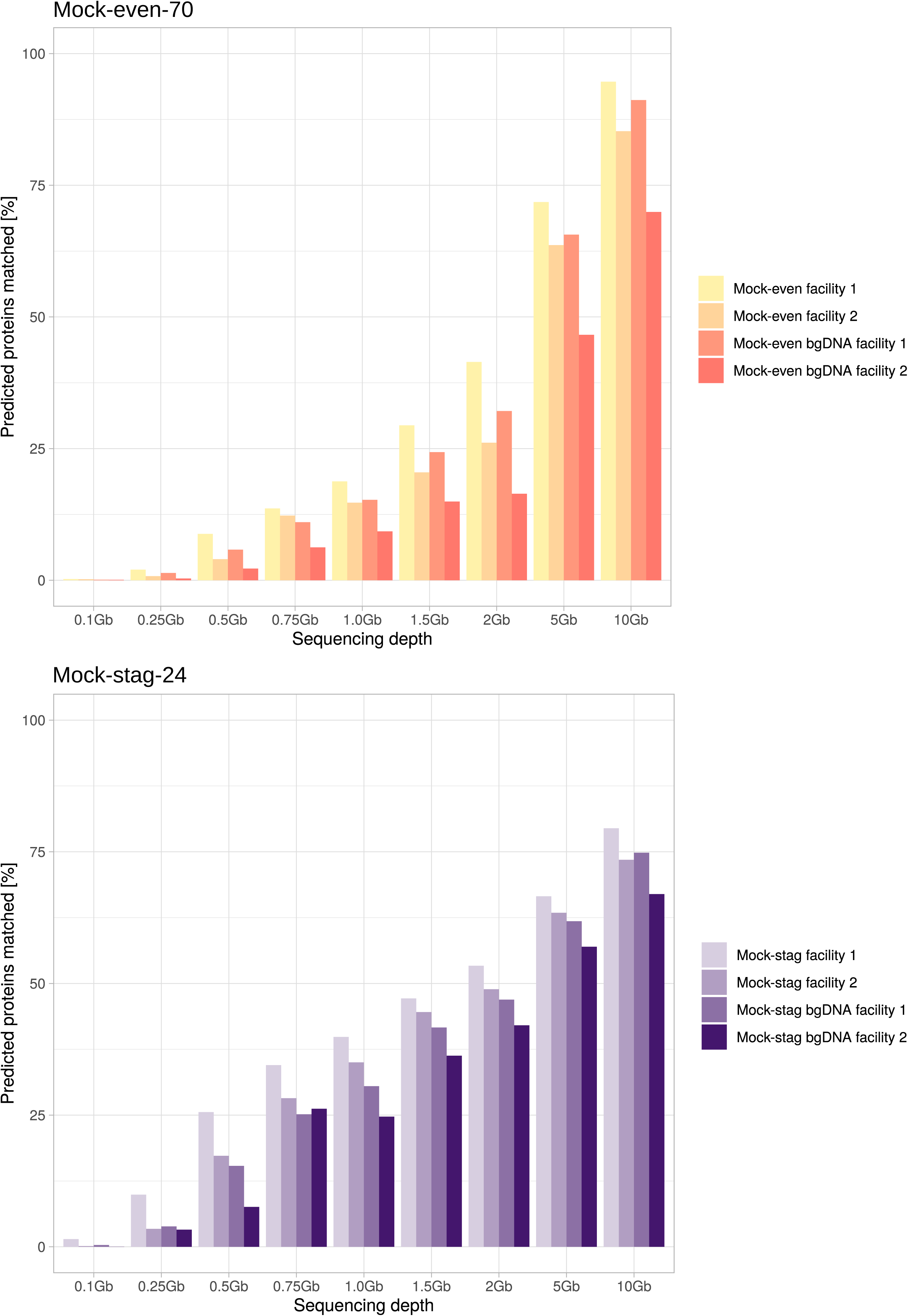
Bar charts of protein sequences recovered. Protein sequenced predicted in the metagenomic datasets as percentage of the protein sequences present in the reference genomes (y-axis). The colour gradient indicates the facility where the library was prepared and the addition of background DNA (bgDNA) from the caecum of germfree mice.

**Supplementary Table S1**: Distribution of the reference genomes DNA in the different mock communities.

**Supplementary Table S2**: Characteristics of the reference genomes.

**Supplementary Table S3**: Number of reads after sequencing and subsampling.

**Supplementary Table S4:** Assignment of the MAGs to reference genomes and using GTDB.

**Supplementary Table S5:** P-values for Fig. 1B, Fig. 1C and Fig. 3C.

## References

1. Klindworth, A. et al. Evaluation of general 16S ribosomal RNA gene PCR primers for classical and next-generation sequencing-based diversity studies. Nucleic Acids Research 41, e1–e1 (2013).

2. Xu, W. et al. Characterization of Shallow Whole-Metagenome Shotgun Sequencing as a High-Accuracy and Low-Cost Method by Complicated Mock Microbiomes. Front. Microbiol. 12, 678319 (2021).

3. Hillmann, B., et al. Evaluating the Information Content of Shallow Shotgun Metagenomics. mSystems 3, 10.1128/msystems.00069-18 (2018).

4. La Reau, A. J. et al. Shallow shotgun sequencing reduces technical variation in microbiome analysis. Sci Rep 13, 7668 (2023).

5. Stothart, M. R., McLoughlin, P. D. & Poissant, J. Shallow shotgun sequencing of the microbiome recapitulates 16S amplicon results and provides functional insights. Molecular Ecology Resources 23, 549–564 (2023).

6. Tremblay, J., Schreiber, L. & Greer, C. W. High-resolution shotgun metagenomics: the more data, the better? Briefings in Bioinformatics 23, bbac443 (2022).

7. Usyk, M. et al. Comprehensive evaluation of shotgun metagenomics, amplicon sequencing, and harmonization of these platforms for epidemiological studies. Cell Reports Methods 3, 100391 (2023).

8. Cattonaro, F., Spadotto, A., Radovic, S. & Marroni, F. Do you cov me? Effect of coverage reduction on metagenome shotgun sequencing studies. F1000Res 7, 1767 (2020).

9. Fritz, A. et al. CAMISIM: simulating metagenomes and microbial communities. Microbiome 7, 17 (2019).

10. Afrizal, A. et al. Enhanced cultured diversity of the mouse gut microbiota enables custom-made synthetic communities. Cell Host & Microbe 30, 1630–1645.e25 (2022).

11. Hitch, T. C. A. et al. Broad diversity of human gut bacteria accessible via a traceable strain deposition system. Preprint at 10.1101/2024.06.20.599854 (2024).

12. Godon, J. J., Zumstein, E., Dabert, P., Habouzit, F. & Moletta, R. Molecular microbial diversity of an anaerobic digestor as determined by small-subunit rDNA sequence analysis. Appl Environ Microbiol 63, 2802–2813 (1997).

13. Asnicar, F. et al. Precise phylogenetic analysis of microbial isolates and genomes from metagenomes using PhyloPhlAn 3.0. Nat Commun 11, 2500 (2020).

14. Cock, P. J. A. et al. Biopython: freely available Python tools for computational molecular biology and bioinformatics. Bioinformatics 25, 1422–1423 (2009).

15. Li, H. bioawk, https://github.com/lh3/bioawk.

16. Li, H. seqtk, https://github.com/lh3/seqtk.

17. Bolger, A. M., Lohse, M. & Usadel, B. Trimmomatic: a flexible trimmer for Illumina sequence data. Bioinformatics 30, 2114–2120 (2014).

18. Bushnell, B. BBMap, sourceforge.net/projects/bbmap/.

19. Li, D., Liu, C.-M., Luo, R., Sadakane, K. & Lam, T.-W. MEGAHIT: an ultra-fast single-node solution for large and complex metagenomics assembly via succinct *de Bruijn* graph. Bioinformatics 31, 1674–1676 (2015).

20. Nurk, S., Meleshko, D., Korobeynikov, A. & Pevzner, P. A. metaSPAdes: a new versatile metagenomic assembler. Genome Res. 27, 824–834 (2017).

21. Aroney, S. T. N., et al. CoverM: Read coverage calculator for metagenomics. Zenodo 10.5281/ZENODO.10531253 (2024).

22. Blanco-Míguez, A. et al. Extending and improving metagenomic taxonomic profiling with uncharacterized species using MetaPhlAn 4. Nat Biotechnol 41, 1633–1644 (2023).

23. Jain, C., Rodriguez-R, L. M., Phillippy, A. M., Konstantinidis, K. T. & Aluru, S. High throughput ANI analysis of 90K prokaryotic genomes reveals clear species boundaries. Nat Commun 9, 5114 (2018).

24. Hyatt, D. et al. Prodigal: prokaryotic gene recognition and translation initiation site identification. BMC Bioinformatics 11, 119 (2010).

25. Aramaki, T. et al. KofamKOALA: KEGG Ortholog assignment based on profile HMM and adaptive score threshold. Bioinformatics 36, 2251–2252 (2020).

26. Graham, E. D., Heidelberg, J. F. & Tully, B. J. Potential for primary productivity in a globally-distributed bacterial phototroph. The ISME Journal 12, 1861–1866 (2018).

27. Buchfink, B., Xie, C. & Huson, D. H. Fast and sensitive protein alignment using DIAMOND. Nat Methods 12, 59–60 (2015).

28. Langmead, B. & Salzberg, S. L. Fast gapped-read alignment with Bowtie 2. Nat Methods 9, 357–359 (2012).

29. Danecek, P. et al. Twelve years of SAMtools and BCFtools. GigaScience 10, giab008 (2021).

30. Kang, D. D. et al. MetaBAT 2: an adaptive binning algorithm for robust and efficient genome reconstruction from metagenome assemblies. PeerJ 7, e7359 (2019).

31. Parks, D. H., Imelfort, M., Skennerton, C. T., Hugenholtz, P. & Tyson, G. W. CheckM: assessing the quality of microbial genomes recovered from isolates, single cells, and metagenomes. Genome Res. 25, 1043–1055 (2015).

32. Chaumeil, P.-A., Mussig, A. J., Hugenholtz, P. & Parks, D. H. GTDB-Tk: a toolkit to classify genomes with the Genome Taxonomy Database. Bioinformatics 36, 1925–1927 (2020).

33. Yu, Y., Ouyang, Y. & Yao, W. shinyCircos: an R/Shiny application for interactive creation of Circos plot. Bioinformatics 34, 1229–1231 (2018).

34. Mattock, J. & Watson, M. A comparison of single-coverage and multi-coverage metagenomic binning reveals extensive hidden contamination. Nat Methods 20, 1170–1173 (2023).

35. De Coster, W. & Rademakers, R. NanoPack2: population-scale evaluation of long-read sequencing data. Bioinformatics 39, btad311 (2023).

36. Hall, M. Rasusa: Randomly subsample sequencing reads to a specified coverage. JOSS 7, 3941 (2022).

37. Kolmogorov, M., Yuan, J., Lin, Y. & Pevzner, P. A. Assembly of long, error-prone reads using repeat graphs. Nat Biotechnol 37, 540–546 (2019).

38. Gurevich, A., Saveliev, V., Vyahhi, N. & Tesler, G. QUAST: quality assessment tool for genome assemblies. Bioinformatics 29, 1072–1075 (2013).

39. R Core Team. R: A Language and Environment for Statistical Computing, https://www.R-project.org/. R Foundation for Statistical Computing, Vienna, Austria https://www.R-project.org/ (2021).

40. Jari Oksanen and Gavin L. Simpson and F. Guillaume Blanchet and Roeland Kindt and Pierre Legendre and Peter R. Minchin and R.B. O’Hara and Peter Solymos and M. Henry H. Stevens and Eduard Szoecs and Helene Wagner and Matt Barbour and Michael Bedward and Ben Bolker and Daniel Borcard and Tuomas Borman and Gustavo Carvalho and Michael Chirico and Miquel {De Caceres} and Sebastien Durand and Heloisa Beatriz Antoniazi Evangelista and Rich FitzJohn and Michael Friendly and Brendan Furneaux and Geoffrey Hannigan and Mark O. Hill and Leo Lahti and Dan McGlinn and Marie-Helene Ouellette and Eduardo {Ribeiro Cunha} and Tyler Smith and Adrian Stier and Cajo J.F. {Ter Braak} and James Weedon. vegan: Community Ecology Package, https://vegandevs.github.io/vegan/. (2024).

41. Hadley Wickham. Flexibly Reshape Data, http://had.co.nz/reshape. (2022).

42. Wickham, H. et al. Welcome to the Tidyverse. JOSS 4, 1686 (2019).

43. Wickham, H., François, R., Henry, L., Müller, K. & Vaughan, D. dplyr: A Grammar of Data Manipulation. 1.1.4 10.32614/CRAN.package.dplyr (2014).

44. Wickham, H. Ggplot2: Elegant Graphics for Data Analysis. (Springer international publishing, Cham, 2016).

45. Alboukadel Kassambara. ggpubr: ‘ggplot2’ Based Publication Ready Plots, https://rpkgs.datanovia.com/ggpubr/. (2023).

46. Xu, S. et al. Use ggbreak to Effectively Utilize Plotting Space to Deal With Large Datasets and Outliers. Front. Genet. 12, 774846 (2021).

47. Yuan Tang and Masaaki Horikoshi. ggfortify: Data Visualization Tools for Statistical Analysis Results, https://CRAN.R-project.org/package=ggfortify. (2016).

48. Meziti, A. et al. The Reliability of Metagenome-Assembled Genomes (MAGs) in Representing Natural Populations: Insights from Comparing MAGs against Isolate Genomes Derived from the Same Fecal Sample. Appl Environ Microbiol 87, e02593–20 (2021).

49. Berg, M., Reiter, T., Emerson, J., Brown, C. T. & Roux, S. Comparison of short-read and long-read metagenome assemblies in a natural soil community highlights systematic bias in recovery of high-diversity populations. NAR Genomics and Bioinformatics 7, lqaf163 (2025).

50. Trigodet, F., Sachdeva, R., Banfield, J. F. & Eren, A. M. Assemblies of long-read metagenomes suffer from diverse errors. Preprint at 10.1101/2025.04.22.649783 (2025).

51. Ma, J. et al. A human gut metagenome-assembled genome catalogue spanning 41 countries supports genome-scale metabolic models. Nat Microbiol 10.1038/s41564-025-02206-1 (2025) doi:10.1038/s41564-025-02206-1.

52. Gweon, H. S. et al. The impact of sequencing depth on the inferred taxonomic composition and AMR gene content of metagenomic samples. Environmental Microbiome 14, 7 (2019).

53. Bowers, R. M. et al. Impact of library preparation protocols and template quantity on the metagenomic reconstruction of a mock microbial community. BMC Genomics 16, 856 (2015).

54. Poulsen, C. S., Ekstrøm, C. T., Aarestrup, F. M. & Pamp, S. J. Library Preparation and Sequencing Platform Introduce Bias in Metagenomic-Based Characterizations of Microbiomes. Microbiol Spectr 10, e00090–22 (2022).

55. Gaulke, C. A. et al. Evaluation of the Effects of Library Preparation Procedure and Sample Characteristics on the Accuracy of Metagenomic Profiles. mSystems 6, 10.1128/msystems.00440-21 (2021).

56. Fierer, N. et al. Guidelines for preventing and reporting contamination in low-biomass microbiome studies. Nat Microbiol 10, 1570–1580 (2025).

57. Meyer, F. et al. Critical Assessment of Metagenome Interpretation: the second round of challenges. Nat Methods 19, 429–440 (2022).

